# Cell wall integrity modulates a PHYTOCHROME-INTERACTING FACTOR (PIF) – HOOKLESS1 (HLS1) signalling module controlling apical hook formation in Arabidopsis

**DOI:** 10.1101/2023.08.05.551077

**Authors:** Riccardo Lorrai, Özer Erguvan, Sara Raggi, Kristoffer Jonsson, Jitka Široká, Danuše Tarkowská, Ondřej Novák, Stéphane Verger, Stéphanie Robert, Simone Ferrari

## Abstract

Etiolated seedlings of dicots form an apical hook to protect the meristems during soil emergence. Hook formation is the result of differential growth on both sides of the hypocotyl apex and is tightly controlled by environmental cues and hormones, among which auxin and gibberellins (GAs) are the main contributors. Cell expansion is tightly regulated by the cell wall, but whether and how feedback from this structure contributes to hook development is still unclear. Here we show that etiolated seedlings of the *Arabidopsis thaliana quasimodo2-1* (*qua2*) mutant, defective in pectin biosynthesis, display severe defects in apical hook formation and maintenance, accompanied by loss of asymmetric auxin maxima and differential cell expansion. Moreover, *qua2* seedlings show reduced expression of *HOOKLESS1* (*HLS1*) and *PHYTOCHROME-INTERACTING FACTOR 4* and *5* (*PIF4/5*), positive regulators of hook formation, and accumulate reduced levels of the active gibberellin GA_4_. Treatment of wild-type seedlings with the cellulose inhibitor isoxaben (isx) also prevents hook development and represses *HLS1* expression and PIF4 accumulation. Moreover, isx stabilizes the DELLA protein REPRESSOR OF *ga1-3* (RGA), which inhibits *HLS1* expression and hook formation. Exogenous GAs or *HLS1* overexpression partially restore hook development in isx-treated seedlings. Notably, agar concentration in the medium restores, both in *qua2* and isx-treated seedlings, hook development and WT-like levels of PIFs and HLS1. We propose that turgor-dependent signals link changes in cell wall integrity to the PIF4/5-HLS1 signalling module to repress differential cell elongation during hook formation.

**Significance statement:** Cell wall integrity modulates apical hook development through poorly understood mechanisms. We show here that, in Arabidopsis, repression of hook formation by either mutations in pectin biosynthesis or by isoxaben treatment is at least partially mediated by the downregulation of a gibberellin-controlled signalling module that comprises PIF4/5 and HLS1. Our results indicate that the signals derived from changes in the cell wall can modulate hormone-mediated pathways to control asymmetric growth during plant development.

## Introduction

Apical hook formation in etiolated seedlings depends on the differential cell elongation on the opposite sides of the hypocotyl apex, causing the shoot to bend by 180° (Guzman and Ecker, 1990; Abbas *et al*., 2013). Like most plant developmental processes, hook formation is largely controlled by the hormone auxin (Abbas *et al*., 2013). Shortly after germination, the formation of an auxin response maximum restrains cell expansion on the concave side of the hook, leading to differential cell elongation and eventually shoot bending (Abbas *et al*., 2013).

In *Arabidopsis thaliana*, hook development is positively controlled by the master regulator HOOKLESS1 (HLS1) (Guzman and Ecker, 1990; Guzman and Ecker, 1990; Lehman *et al*., 1996; Li *et al*., 2004; Zhang *et al*., 2018). HLS1 was reported to promote the asymmetric distribution of auxin between the concave and convex sides of the hypocotyl (Lehman *et al*., 1996) and to reduce the levels of AUXIN RESPONSE FACTOR 2 (ARF2), a repressor of auxin responses (Li *et al*., 2004). Both apical hook formation and *HLS1* expression are promoted by ethylene and gibberellins (GAs) (Lehman *et al*., 1996; An *et al*., 2012) and negatively regulated by jasmonates (Song *et al*., 2014). Regulation of hook development by GAs is mediated by the degradation of the key repressors DELLA proteins (Sun, 2008). When GA levels are low, DELLAs promote the proteasome-mediated degradation of PHYTOCHROME-INTERACTING FACTORS (PIFs) (Li *et al*., 2016), a family of transcription factors that positively regulate the expression of *HLS1* (Zhang *et al*., 2018). In addition, DELLAs inhibit the activity of PIFs by sequestering their DNA-recognition domain (De Lucas *et al*., 2008; Feng *et al*., 2008). On the other hand, jasmonates can repress hook formation by reducing *HLS1* expression (Zhang *et al*., 2014) and by repressing PIF function (Zhang *et al*., 2018).

Increasing evidence indicates that the hormone-mediated signalling pathways that modulate hook formation are modulated by the cell wall (Aryal *et al*., 2020; Baral *et al*., 2021; Jonsson *et al*., 2021). Primary cell walls are mainly composed of cellulose, hemicelluloses, and pectin, which determine their mechanical properties, like resistance to tensile stress, thus regulating cell shape and organ morphogenesis (Cosgrove, 2005). A correlation between pectin composition, auxin responses and differential cell elongation during hook development was recently reported (Jonsson *et al*., 2021). Cells in the inner side of the hook, where auxin maxima occur, display a higher degree of methylesterification (DM) of homogalacturonan (HG), a major pectic polysaccharide, which correlates with a reduction in cell elongation (Jonsson *et al*., 2021). Notably, plants overexpressing a pectin methylesterase inhibitor show a uniformly high pectin DM and fail to establish a proper auxin response gradient, resulting in defective hook formation (Jonsson *et al*., 2021). These observations suggest that pectin composition and/or architecture directly modulates the signalling pathways that control differential cell expansion in the hypocotyl. In addition, alterations in other cell wall structural components, including cellulose (Sinclair *et al*., 2017; Baral *et al*., 2021) and xyloglucan (Aryal *et al*., 2020) also impair apical hook development, suggesting that changes in different wall structural components converge into common responses that restrict differential cell elongation, preventing hook formation. However, the exact mechanisms linking cell wall composition to the signalling events that regulate hook development are not fully elucidated.

Plants have evolved mechanisms to monitor cell wall integrity (CWI) and, in case of alterations, mount compensatory responses that reinforce the cell wall to ensure its functional integrity and that can restrict cell expansion (Ellis *et al*., 2002; Engelsdorf *et al*., 2018). CWI can be experimentally altered by lesions in genes involved in the biosynthesis of cell wall structural polysaccharides or by chemicals that interfere with it, like isoxaben (isx), an inhibitor of cellulose deposition (Vaahtera *et al*., 2019). Etiolated Arabidopsis seedlings with altered cellulose deposition display strongly reduced hypocotyl growth (Fagard *et al*., 2000) and accumulate high levels of jasmonates (Engelsdorf *et al*., 2018). Defects in pectin composition also restrict the growth of etiolated hypocotyls. Two Arabidopsis mutants impaired in HG biosynthesis, namely *QUASIMODO1* (*QUA1*), encoding a putative glycosyltransferase (Bouton *et al*., 2002), and *QUA2*/*TUMOROUS SHOOT DEVELOPMENT 2* (*TSD2*), encoding a Golgi-localized pectin methyltransferase (Krupková *et al*., 2007; Mouille *et al*., 2007; Du *et al*., 2020), have shorter hypocotyls and defects in hypocotyl epidermis cell elongation and adhesion (Krupková *et al*., 2007; Mouille *et al*., 2007; Raggi *et al*., 2015). Turgor-sensitive processes appear to be relevant for the detection of CWI changes and the activation of downstream responses that restrict growth. For instance, several responses induced by isx are largely sensitive to osmotic manipulation, *i.e.*, co-treatments with osmotic (Hamann *et al*., 2009; Engelsdorf *et al*., 2018). Similarly, cell adhesion and elongation defects in *qua1* are suppressed by reducing external water potential *via* increased agar concentration in the growth medium (Verger *et al*., 2018). Whether and how these responses can also affect developmental processes regulated by differential cell elongation, as in the case of apical hook formation, is currently unclear. Here we report that loss of CWI represses a signalling module that comprises PIF4/5 and HLS1, resulting in a defective apical hook, and that these effects are suppressed by high agar concentration of the growth medium. Our results suggest that turgor-dependent responses to altered CWI directly modulate signalling events that control differential cell expansion during hook development.

## Results

### Defects in pectin biosynthesis impair hook formation and maintenance in a turgor-dependent manner

Apical hook formation was initially examined in a panel of *Arabidopsis* mutants impaired in different cell wall polysaccharides to determine the relative impact of changes in specific cell wall components on this process. Under our experimental conditions, four-day-old etiolated wild-type (WT) seedlings displayed a completely closed hook (Figure 1a-b), which, in contrast, was completely open in *qua2* as well as in two other mutants affected in pectin composition, *gae1 gae6* and *murus1* (*mur1*) (Fig 1a-b). The *gae1 gae6* double mutant carries mutations in two glucuronate 4-epimerases (GAEs) required for the biosynthesis of UDP-D-galacturonic acid (Mølhøj *et al*., 2004) and is defective in HG (like *qua2*) and, possibly, rhamnogalacturonan I (RG-I) biosynthesis (Bethke *et al*., 2016), while *mur1* is impaired in fucose biosynthesis (Bonin *et al*., 1997) and has therefore defective RG-II, xyloglucans and cell wall glycoproteins (Reiter *et al*., 1993; Rayon *et al*., 1999; Freshour *et al*., 2003).

**Figure 1.**
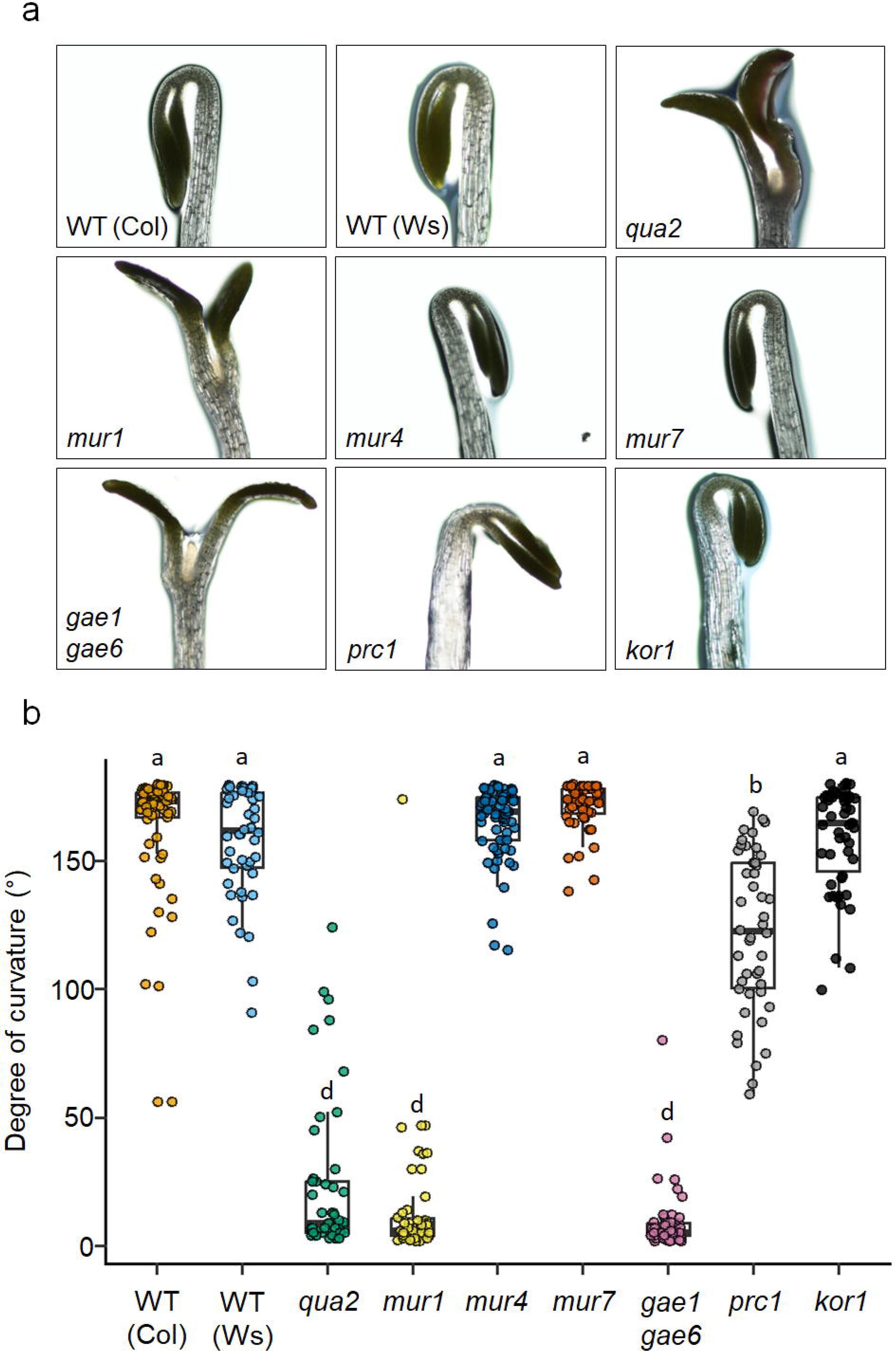
Development of apical hook in Arabidopsis cell wall mutants. (**a**) Representative pictures of wild-type (WT) Columbia-0 (Col), WT Wassilewskija (Ws), *qua2*, *mur1*, *mur4*, *mur7*, *gae1gae6*, *prc1* (in Col-0 background) and *kor1* (in Ws background) four-day-old dark-grown seedlings. Scale bar, 0.5 mm. (**b**) Quantification of apical hook angles of seedlings grown as in (**a**). Box plots indicate the 1^st^ and 3^rd^ quartiles split by median; whiskers show range (n>20). Letters indicate statistically significant differences (P < 0.05) according to one-way ANOVA followed by post-hoc Tukey’s HSD.

We also examined *procuste1* (*prc1*) and *korrigan1* (*kor1*), carrying mutations in a cellulose synthase subunit and in an endo-1,4-β-glucanase, respectively, and therefore impaired in cellulose deposition in primary cell walls (Desnos *et al*., 1996; Nicol *et al*., 1998). *prc1* showed only a mild defect in hook formation, whereas *kor1* was comparable to the wild type (Figure 1a-b). No significant defects in hook development were observed in *mur4* and *mur7* seedlings (Figure 1a-b), both impaired in the biosynthesis of arabinose (Reiter *et al*., 1997; Burget *et al*., 2003), a constituent of RG-I, RG-II, hemicelluloses and glycoproteins such as arabinogalactan proteins (Kaczmarska *et al*., 2022). Taken together, these results suggest that mutations in genes regulating HG biosynthesis have a major impact on hook formation, compared to defects in other wall components.

Turgor pressure mediates the activation of several responses triggered by altered CWI (Hamann *et al*., 2009; Engelsdorf *et al*., 2018). To verify if turgor-dependent responses also mediate the effects of altered pectin composition on hook formation and to determine what phases of this process are specifically affected by the mutations, a kinematic analysis of hook development was performed in WT, *qua2*, *gae1gae6* and *mur1* seedlings grown in the dark on medium containing 0.8% (w/v) or 2.5% (w/v) agar [henceforth indicated as low agar (LA) and high agar (HA), respectively]. WT seedlings grown on LA displayed a typical (Abbas *et al*., 2013) formation, maintenance and opening phase (Figure 2a-c). In contrast, all mutants grown on LA showed a formation phase comparable, in length, to the WT, but were unable to form a fully closed hook (Figure 2a-c). Moreover, the maintenance phase was deeply compromised in all mutants, leading to hook opening right after the maximum curvature was achieved (Figure 2a-c).

**Figure 2.**
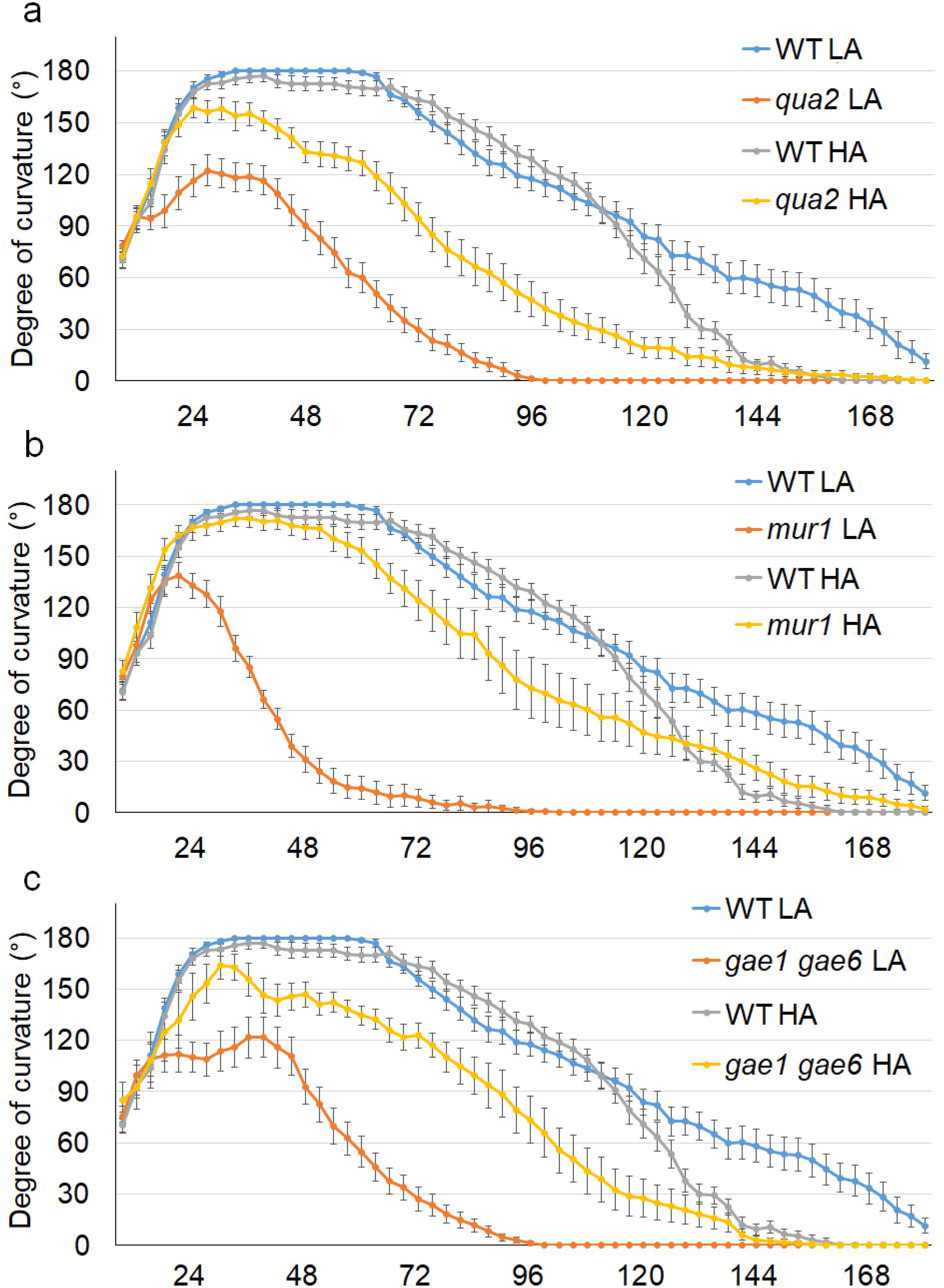
Kinematic analysis of apical hook formation in pectin mutants grown onlow and high agar. Wild-type (WT, blue and grey lines) and *qua2* (a), *mur1* (b) or *gae1gae6* (c) mutant (orange and yellow lines) seedlings were grown in the dark on medium containing either 0.8% (w/v) (LA, blue and orange lines) or 2.5% (w/v) agar (HA, gray and yellow lines). The hook angle was measured at the indicated times. Error bars represent mean angle ± SE (n≥15).

When WT seedlings were grown on HA, formation and maintenance of the hook were largely unaffected, though the opening phase was accelerated (Figure 2a-c). Notably, growth on HA partially restored hook formation in all mutant lines (Figure 2a-c), leading to a statistically significant increase of the maximum angle of curvature (Figure S1). In addition, HA also rescued the maintenance phase in *mur1* seedlings (Figure 2c). Hook development could also be restored by sorbitol, an osmolyte previously shown to suppress responses induced by cell wall damage (Hamann *et al*., 2009; Engelsdorf *et al*., 2018) (Figure S2). Taken together, these results suggest that turgor-dependent responses are responsible for the impaired hook development observed in seedlings with altered pectin composition.

### Loss of pectin integrity disrupts differential cell expansion and asymmetric auxin response during apical hook development

Hook formation is largely dependent on the differential elongation rate of epidermal cell on the two sides of the hypocotyl (Lehman *et al*., 1996). Defects in *QUA2* restrict cell expansion in the epidermis of adult leaves (Raggi *et al*., 2015), suggesting that alterations in cell expansion rates might also occur in the epidermis of the hypocotyl of *qua2* etiolated seedlings, resulting in a defective hook. Individual cell elongation rates were therefore measured in the apical portion of the hook of WT and *qua2* seedlings grown in the dark in LA and HA condition. As expected, cell expansion rate in WT seedlings was lower in the inner side than in the outer side of the hypocotyl, either in LA or HA condition (Figure 3a-b). In contrast, *qua2* seedlings showed a significant reduction in the expansion rate in the outer side of the hook when grown on LA, but not on HA (Figure 3a-b).

**Figure 3.**
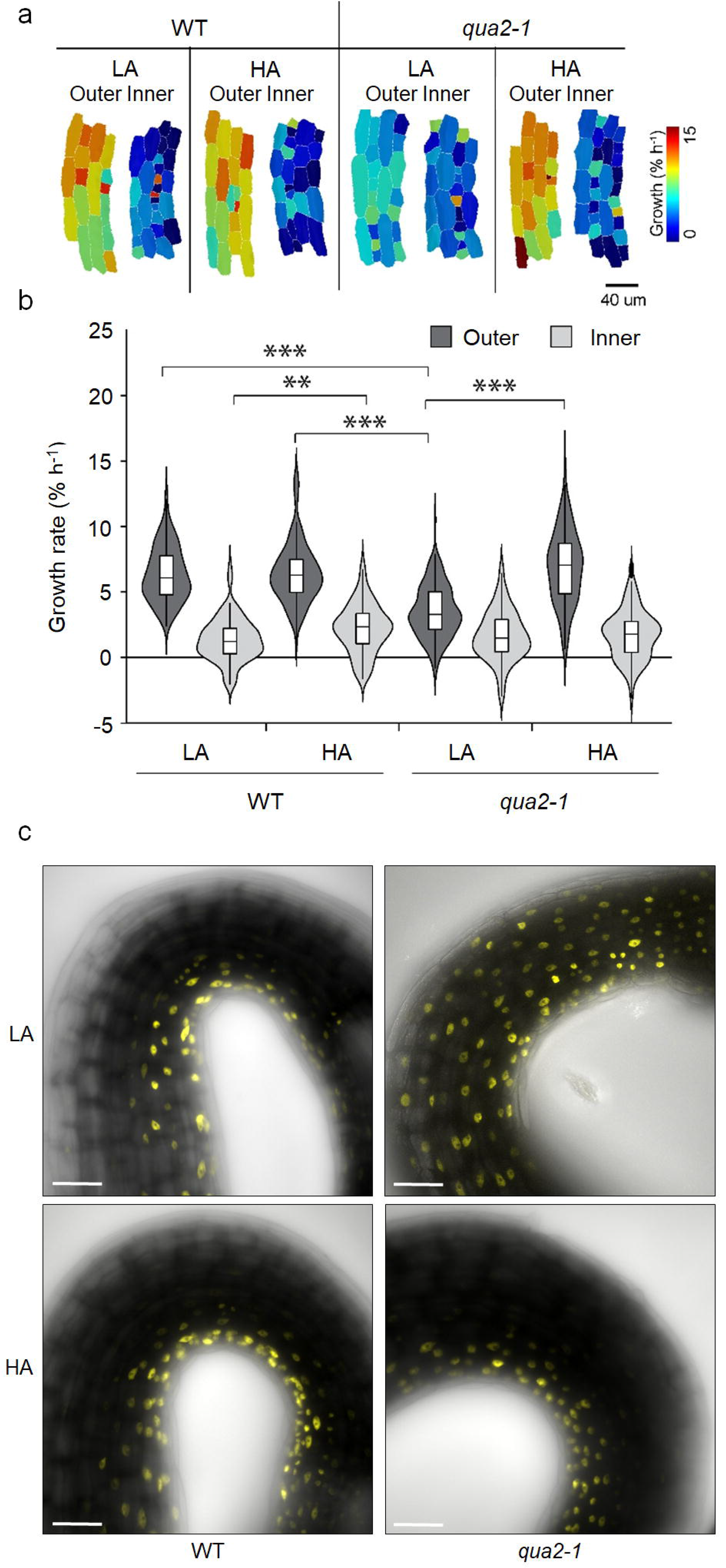
Effects of agar concentration on cell elongation and auxin response during apical hook formation in *qua2* seedlings. (**a**) Heatmaps of growth rate of individual cells in the apical portion of the hypocotyl upon three-hour time lapse in wild-type (WT) and *qua2* seedlings grown in the dark on medium containing 0.8% (LA) or 2.5% (LA) (w/v) agar. (**b**) Quantification of growth rate of individual cells in the outer (dark gray) and inner (light gray) side of the hypocotyl of seedlings grown as in (**a**). Data are average of three independent biological replicates ± SD. For each experiment, 15 cells from both inner and outer side of the hook were measured from each of 9 individual seedlings. Asterisks indicate statistical significance by Student’s t test (**, p < 0.01; ***, p < 0.001). (**c**) Representative confocal laser scanning microscopy images of WT and *qua2* seedlings expressing the DR5::Venus-NLS and grown in the dark on LA or HA. Scale bars, 50 μm.

As differential cell expansion is dependent on the establishment of an auxin gradient at the two sides of the apex (Abbas *et al*., 2013), the distribution of auxin signalling was evaluated in WT and *qua2* seedlings expressing the auxin response marker *DR5*-VENUS-NLS (Heisler *et al*., 2005). WT seedlings displayed a strong fluorescent signal predominantly in the inner epidermal cells of the hook and this pattern was not affected by the agar concentration in the medium (Figure 3c). In contrast, reporter expression was equally distributed on both sides of the hypocotyl of *qua2* seedlings grown in LA (Figure 3c). This alteration was fully restored when the mutant was grown on HA (Figure 3c). Taken together, our results indicate that turgor-dependent responses to altered HG hinder proper asymmetric auxin signalling and differential cell expansion during hook formation.

### Loss of pectin integrity represses *HLS1* and *PIF* expression and GA accumulation in dark-grown seedlings

HLS1 combines upstream stimuli important for hook formation (Guzman and Ecker, 1990), negatively regulating ARF2 levels (Li *et al*., 2004) and influencing auxin distribution (Lehman *et al*., 1996). Hook formation is also positively modulated by PIFs, and in particular PIF4 directly binds to the promoter of *HLS1* to activate its transcription (Zhang *et al*., 2018). We therefore evaluated if a defective pectin composition might affect the expression of the genes encoding these proteins. Indeed, transcript levels for both *HLS1* and *PIF4* were significantly reduced in etiolated *qua2*, *mur1* or *gae1 gae6* seedlings grown in LA, in comparison to the wild type (Figure 4a-b). Expression of both genes in the mutants was restored by HA, which indeed increased *HLS1* transcript levels also in the WT (Figure 4a). The expression of *PIF5*, previously shown to positively regulate hook maintenance (Khanna *et al*., 2007), was also reduced in *qua2* seedlings grown on LA, but not on HA (Figure 4c).

**Figure 4.**
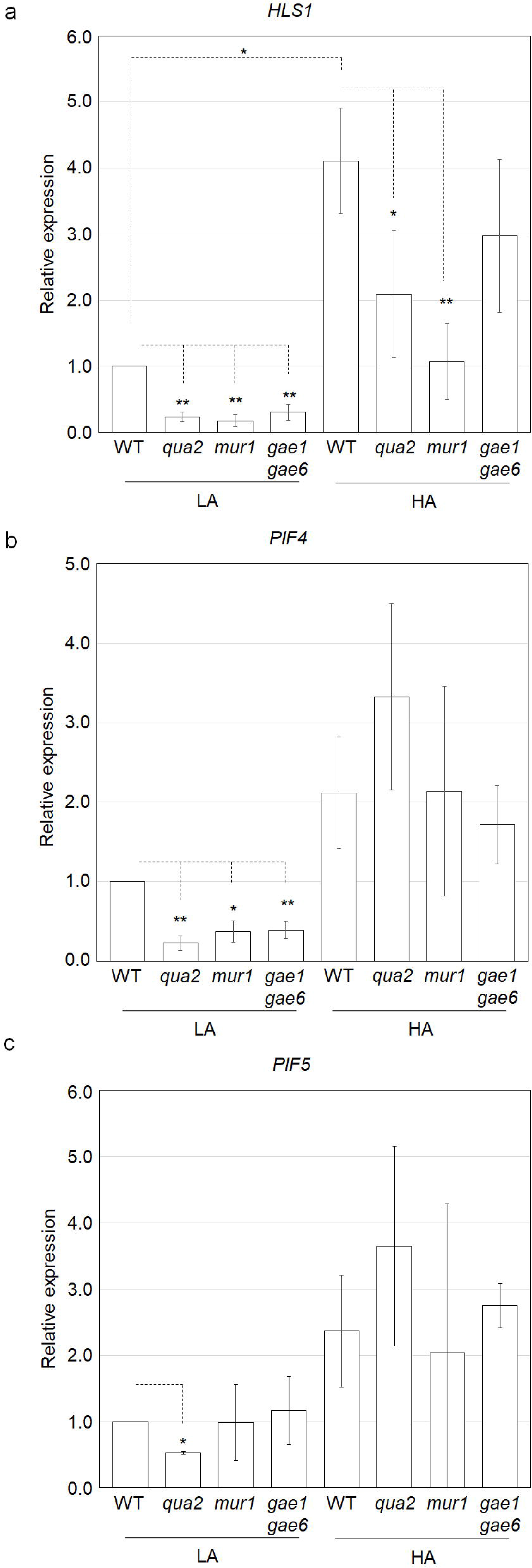
*HLS1* and *PIF4/5* expression in pectin mutants. Total RNA was extracted from three-day-old wild-type (WT), *qua2*, *mur1*, and *gae1gae6* seedlings grown in the dark on medium containing 0.8% (LA) or 2.5% (HA) agar (w/v). Relative transcript levels of *HLS1* (**a**), *PIF4* (**b**) and *PIF5* (**c**) were measured by qPCR, using *UBQ5* as reference. Data are means of n≥3 independent biological replicates ± SD. Asterisks indicate statistical significance according to Student’s t test (*P < 0.05; **P < 0.01).

Hook formation and *HLS1* expression are both positively regulated by GAs (An *et al*., 2012). To evaluate if loss of pectin integrity affects the levels of these hormones in a turgor-dependent manner, levels of GA_4_, the major active GA in Arabidopsis seedlings (Yamaguchi, 2006), were quantified in WT and *qua2* etiolated seedlings grown in the dark on LA or HA medium. Under LA conditions, GA_4_ levels were significantly lower in the mutant, compared to the wild type (Figure 5a). In contrast, GA_4_ levels in *qua2* seedlings grown on HA were like those observed in the wild type grown on LA (Figure 5a). Unexpectedly, GA_4_ levels were significantly reduced in WT seedlings grown on HA (Figure 5a). Consistent with the observed reduction in GA_4_ accumulation, transcript levels for *GA20ox1* and *GA3ox1*, required for GA_4_ biosynthesis (Hedden and Phillips, 2000), were reduced in *qua2* and, to a lesser extent, *gae1gae6* and *mur1* etiolated seedlings grown on LA (Figure 5b-c). Notably, transcript levels of *GA20ox1* and *GA3ox1* significantly increased in both WT and mutant seedlings grown in HA (Figure 5b-c).

**Figure 5.**
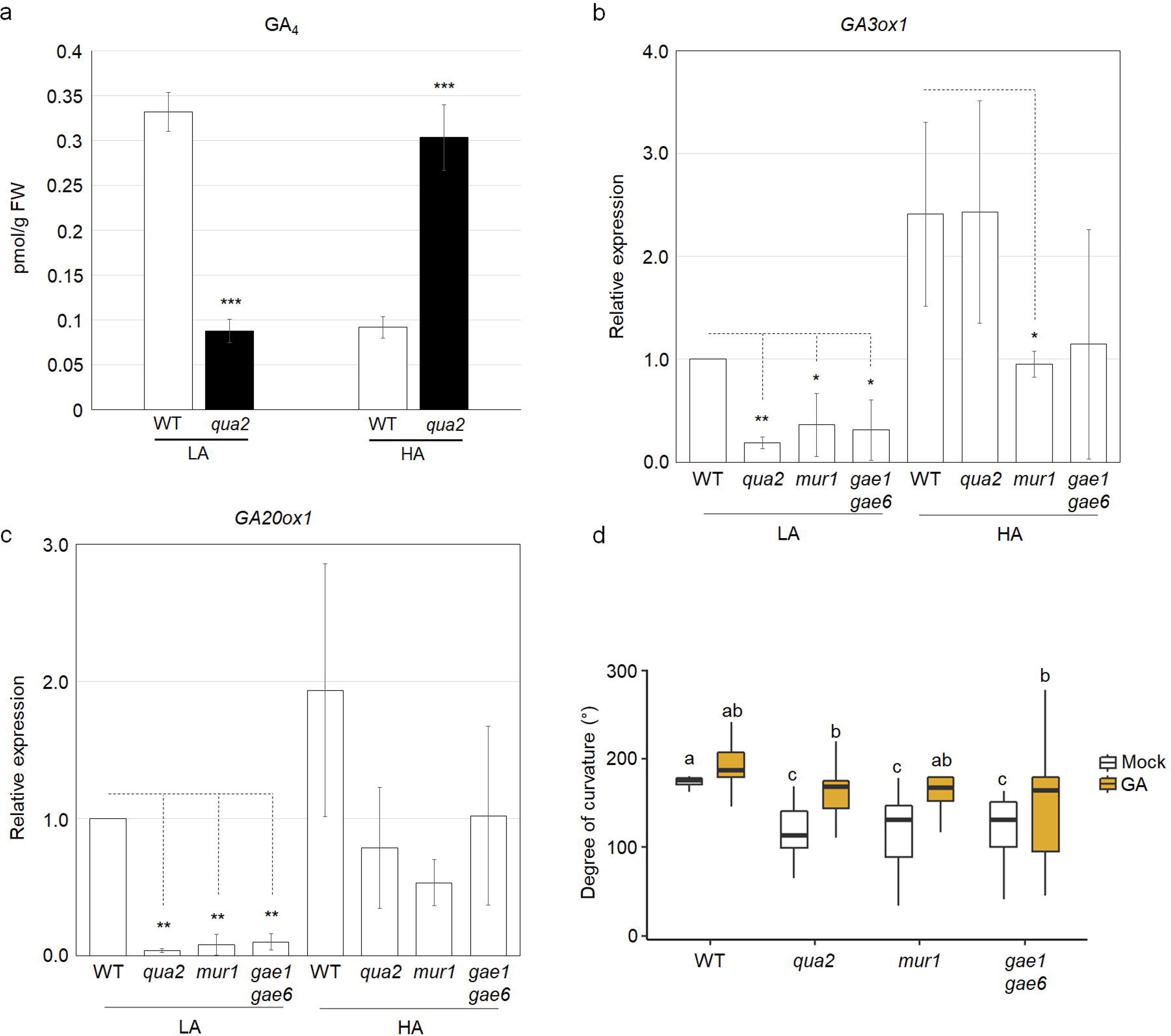
Defects in GA accumulation and expression of biosynthetic genes in pectin mutants. (**a**) Levels of the bioactive GA_4_ in three-day-old WT (white bars) and *qua2* (black bars) seedlings grown in the dark on medium containing 0.8% (w/v) or 2.5% (w/v) agar (LA and HA, respectively). Bars represent the mean of three independent biological replicates ± SD. Asterisks indicate statistically significant differences between WT and *qua2*, according to Student’s t-test (*, p≤0.05; **, p≤0.01; ***, p≤0.001). (**b**, **c**) Expression of *GA3ox1* (**b**) and *GA20ox1* (**c**) in WT, *qua2*, *mur1* and *gae1gae6* seedlings grown as in (**a**). Transcript levels were determined by qPCR using *UBQ5* as reference. Bars indicate mean relative expression, compared to WT seedlings grown in LA, ± SD of at least three independent biological replicates. Asterisks indicate statistically significant differences with WT in LA conditions, according to Student’s t-test (*, p≤0.05; **, p≤0.01). (**d**) Apical hook angles of three-day-old WT, *qua2*, *mur1* and *gae1gae6* seedlings grown in the dark on medium supplemented with ethanol (mock, white boxes) or 50 μM GA_4_ (GA, yellow boxes). Box plots indicate the 1^st^ and 3^rd^ quartiles split by median; whiskers show range. Letters indicate statistically significant differences, according to two-way ANOVA followed by post-hoc Tukey’s HSD (p< 0.05).

To evaluate if reduced levels of GAs might contribute to the altered hook formation observed in pectin-related mutants, apical hook angle was measured in WT, *qua2*, *gae1gae6* and *mur1* seedlings grown on LA in the absence or presence of exogenous GAs. Indeed, exogenous GAs restored almost WT-like hook formation in all mutants (Figure 5d). Taken together, these results suggest that responses triggered by loss of pectin integrity, and dependent on turgor pressure, lead to a reduction of the levels of active GAs in etiolated seedlings, possibly *via* the repression of GA biosynthetic genes, and that such reduced GA levels might contribute to repress downstream signalling events that promote hook formation.

### Isoxaben inhibits hook formation and represses *HLS1* and *PIF4/5* expression in a turgor-dependent manner

To investigate if turgor-dependent repression of the PIFs/HLS1 signalling module is a specific response to altered pectin composition or a general response to loss of CWI, a pharmacological approach was adopted, growing etiolated WT seedlings in the presence of increasing concentrations of isx. Under LA conditions, isx compromised hook curvature at concentrations equal to or higher than 2.5 nM (Figure 6a). HA conditions restored hook formation in the presence of isx at a dose of 2.5 nM and, to a lesser extent, 5.0 nM (Figure 6a). As in pectin mutants, isx repressed the expression of *GA3ox1* and *GA20ox1*, and this repression was alleviated by HA (Figure S3a-b). Moreover, expression of the GA2-oxidase gene *GA2ox2*, involved in GA_4_ catabolism (Hedden and Phillips, 2000), increased in response to isx in LA, but not in HA conditions (Figure S3c). Isx also reduced transcript levels of *HLS1, PIF4* and *PIF5* under LA, but not HA conditions (Figure 6b-c; Figure S4a). To further investigate the role of *HLS1* down-regulation in the repression of hook development induced by isx, hook curvature was measured in transgenic *hls1* seedlings overexpressing myc-HLS1 (Shen *et al*., 2016). The overexpression of HLS1 fully restored hook development in the presence of 2.5 nM isx (Figure 6d), though at higher doses hook formation was still compromised (Figure 6d). These results suggest that reduced *HLS1* expression contributes to the defective hook formation caused by loss of CWI.

**Figure 6.**
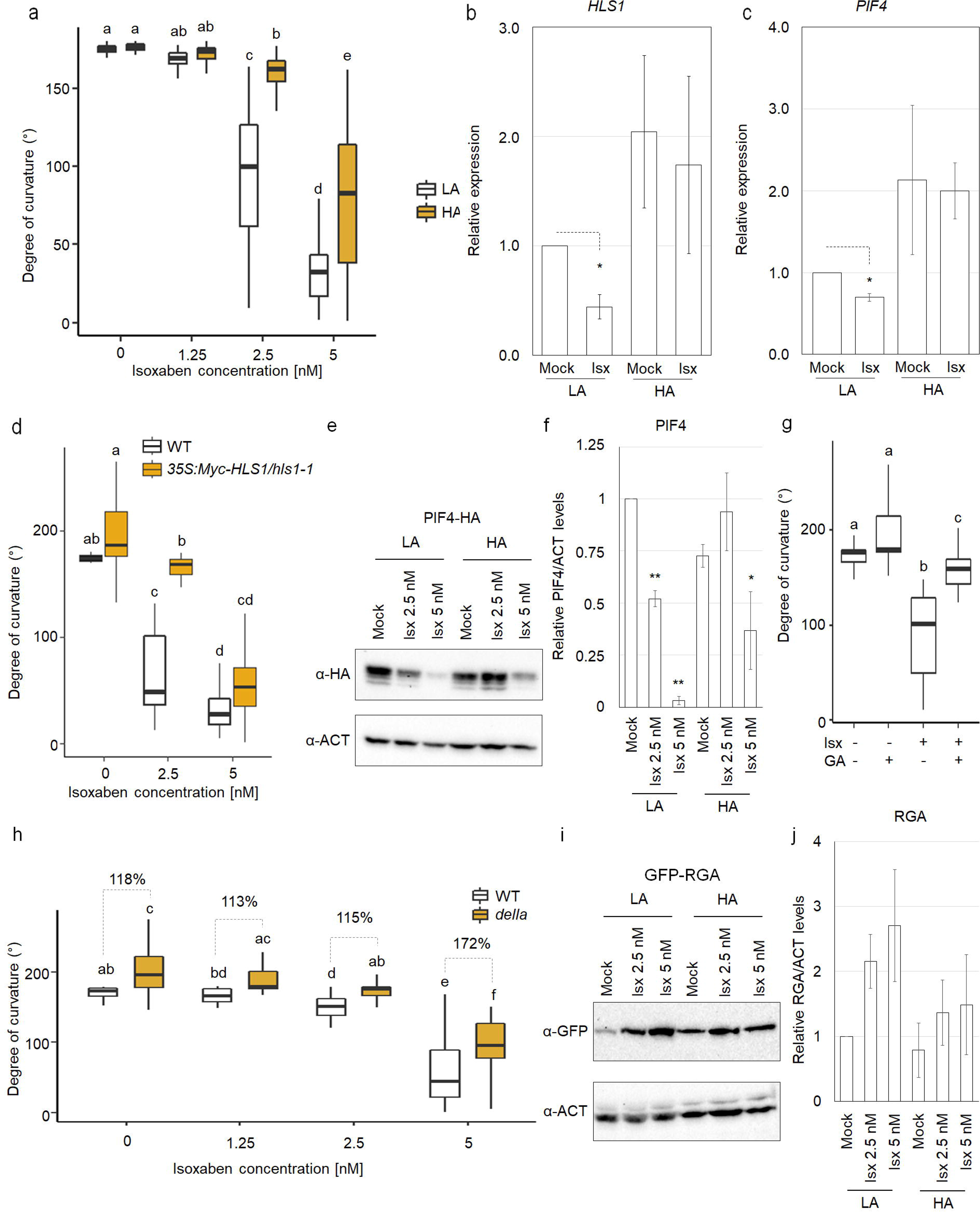
Isoxaben inhibits apical hook formation and GA-mediated signalling in a turgor-dependent manner. (**a**) Quantification of apical hook angles of three-day-old wild-type (WT) seedlings grown in the dark on medium 0.8% (LA) or 2.5% (HA) agar (w/v) and supplemented with isoxaben (isx) at the indicated doses. (**b**, **c**) Transcript levels of *HLS1* (**b**) and *PIF4* (**c**) in three-day-old WT seedlings grown as in (**a**). Transcript levels were determined by qPCR using *UBQ5* as reference gene. Bars indicate mean relative expression levels, compared to WT seedlings grown in LA in the absence of isx, ± SD of at least three independent biological replicates. Asterisks indicate statistically significant differences with WT in LA conditions, according to Student’s t-test (*, p≤0.05; **, p≤0.01). (**d**) Quantification of apical hook angles of wild-type (WT) and 35S:Myc-HLS1/*hls1-1* three-day-old seedlings grown in the dark in the presence of the indicated concentrations of isx (WT, white boxes; 35S:Myc-HLS1/*hls1-1*, yellow boxes). (**e**, **f**) Transgenic lines expressing PIF4-HA under the control of its native promoter (ProPIF4:PIF4-3×HA) were grown on LA or HA medium supplemented with the indicated concentrations of isx. PIF4-HA levels were detected by immunoblot analysis with an antibody against HA; an antibody against actin (ACT) was used as loading control. Bars in (**f**) indicate mean relative normalized intensity ± SD of the PIF4-HA signal, compared to mock-treated seedlings under LA condition (n≥3 independent biological replicates). Asterisks indicate statistically significant differences between mock and isx-treated seedlings at the same agar concentration, according to Student’s t test (*P < 0.05; **P < 0.01). (**g**) Apical hook angles of three-day-old dark-grown WT seedlings treated with DMSO or 2.5 nM isoxaben (isx) in the presence or absence of 50 μM GAs. (**h**) Apical hook angles of wild-type Ler (WT, white boxes) and *della* (yellow boxes) three-day-old seedlings grown in the dark in the presence of isx at the indicated doses. (**i**, **j**) Transgenic lines expressing GFP-RGA under the control of their native promoters were germinated and grown in the dark for three days on LA or HA medium supplemented with DMSO (mock) or isx at the indicated doses. GFP-RGA levels were detected by immunoblot analysis with an antibody against GFP. Intensity of the GFP-RGA band normalized for the loading control ACT and expressed as relative levels compared to mock-treated seedlings under LA condition. Bars in **(j)** indicate mean relative levels ± SD (n≥3 independent biological replicates). Letters in (**a)**, **(d)**, **(g)** and **(h)** indicate statistically significant differences according to two-way ANOVA followed by post-hoc Tukey’s HSD (p<0.05). Box plots indicate the 1^st^ and 3^rd^ quartiles split by median; whiskers show range (n>20).

To determine if PIFs protein levels are significantly affected by isx, transgenic lines expressing a HA-tagged version of PIF4 or PIF5 (de Wit *et al*., 2016; Zhang *et al*., 2017) were grown in the dark on LA or HA medium. A significant reduction of PIF4-HA abundance was observed in seedlings grown on LA in the presence of isx, whereas the impact of isx was much less evident when seedlings were grown on HA and was significant only at the highest dose (Figure 6e-f). PIF5-HA levels were also repressed by high isx doses, when seedlings were grown in LA, but not in HA conditions (Figure S4b-c). As with pectin mutants (Figure 5d), exogenous GAs partially restored hook formation in seedlings treated with isx (Figure 6g), further supporting the hypothesis that suppression of GA-mediated signalling might contribute to the defective hook observed in response to loss of CWI. Since PIF4 stability and *HLS1* expression are negatively regulated by DELLA proteins (An *et al*., 2012; Li *et al*., 2016), we examined the effects of isx on hook curvature in a pentuple mutant (*della*) for all *DELLA* genes (Feng *et al*., 2008). Even though hook curvature was still significantly reduced, the *della* line was less sensitive to isx (Figure 6h). We then evaluated the effects of isx treatment on DELLA protein levels in a transgenic line expressing GFP-RGA (Silverstone *et al*., 2001). GFP-RGA levels significantly increased in response to isx in seedlings grown in LA, but not in HA conditions (Figure 6i-j). Taken together, these results indicate that, as in the case of mutations that impair pectin integrity, isx treatments repress GA-dependent signalling events that control *PIF4/5* and *HLS1* expression and hook curvature and suggest a common mechanism underlying the effects of loss of CWI caused by alterations in different cell wall components on hook development. Moreover, inhibition of hook formation in plants with altered CWI is accompanied by a stabilisation of DELLA proteins, which might be a consequence of reduced GA levels.

### Jasmonates are not involved in defective hook formation caused by altered cell wall integrity

Isx induces the accumulation of jasmonates in *Arabidopsis* seedlings (Engelsdorf *et al*., 2018). In etiolated seedlings, exogenous jasmonic acid (JA) antagonises apical hook development (Song *et al*., 2014; Zhang *et al*., 2014). We reasoned that the altered apical hook development observed in response to loss of CWI might be mediated by increased jasmonate levels. Levels of JA, jasmonyl-L-isoleucine (JA-Ile) and of the JA-derivative 11– and 12-hydroxyjasmonate (Σ 11-/ 12-OHJA, sum of unresolved 11– and 12-OHJA) were therefore quantified in dark-grown WT and *qua2* seedlings. Under LA conditions, mutant seedlings contained higher levels of all three jasmonates, compared to the wild type (Figure 7a). Under HA conditions, the concentration of JA in WT seedlings was unaltered, while JA-Ile and Σ 11-/ 12-OHJA levels were moderately increased (Figure 7a). Growth on HA medium significantly reduced JA and JA-Ile levels in *qua2*, while Σ 11-/ 12-OHJA concentration in the mutant was slightly increased (Figure 7a).

**Figure 7.**
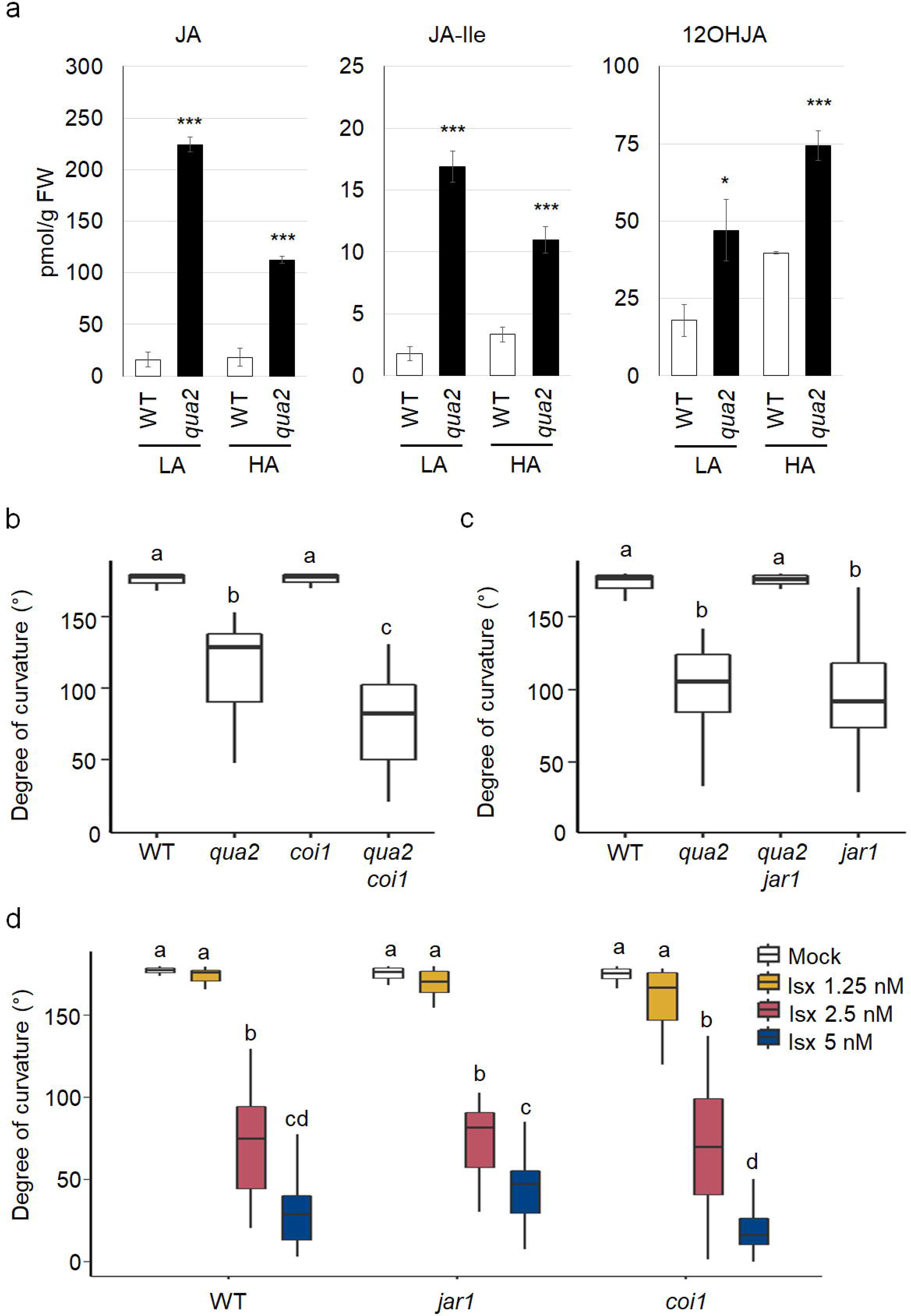
Inhibition of apical hook formation in response to altered cell wall integrity is independent of jasmonate signalling. (a) Levels of JA, JA-Ile, Σ 11-/ 12-OHJA in three-day-old wild-type (WT, white bars) and *qua2* (black bars) seedlings grown in the dark on medium containing 0.8% (LA) or 2.5% (HA) agar (w/v). Bars represent means of three independent biological replicates ± SD. Asterisks indicate significant differences relative to WT, according to Student’s t-test (*p≤0.05, **p≤0.01, ***p≤0.001). (b-c) Apical hook angles of WT, *qua2*, *coi1* and *qua2 coi1* (b), or *jar1* and *qua2 jar1* (c) grown as in (a). (d) Quantification of apical hook angles of wild type (WT Columbia), *jar1* and *coi1* three-day-old seedlings grown in the dark in the presence of isoxaben (isx) at the indicated doses (mock, white boxes; 1.25 nM isx, yellow boxes; 2.5 nM isx, red boxes; 5 nM isx, blue boxes). Box plots in (b-d) indicate the 1^st^ and 3^rd^ quartiles split by median, and whiskers show range. Letters indicate statistically significant differences (p<0.05) according to two-way ANOVA followed by post-hoc Tukey’s HSD.

To assess whether high jasmonates levels are causing hook development alteration of *qua2*, *qua2* was crossed with lines defective for *JASMONATE RESISTANT 1* (*JAR1*), required for the synthesis of JA-Ile (Wasternack and Hause, 2013), or *CORONATINE INSENSITIVE1* (*COI1*), a crucial component of the SCF COI1 E3 ubiquitin complex necessary for JA-Ile perception and transduction (Wasternack and Hause, 2013). In *qua2 coi1* seedlings, hook impairment was slightly exacerbated (Figure 7b), while the *qua2 jar1* double mutant did not show differences in hook angle, compared to *qua2* (Figure 7c). Consistently, *jar1* and *coi1* single mutants treated with isx displayed hook defects comparable to those observed in the wild type (Figure 7d). These results indicate that, despite loss of CWI triggers the accumulation of elevated levels of jasmonates in a turgor-dependent manner, these hormones do not contribute to the observed defects in hook formation.

Taken together, our results suggest that loss of CWI triggers turgor-dependent responses that suppress GA accumulation and GA-mediated downstream signalling events, including *PIF* and *HLS1* expression that positively regulate hook formation (Figure 8). These effects are independent of jasmonate-mediated signalling, and likely disrupt auxin response asymmetry, differential cell elongation and proper hook formation.

**Figure 8.**
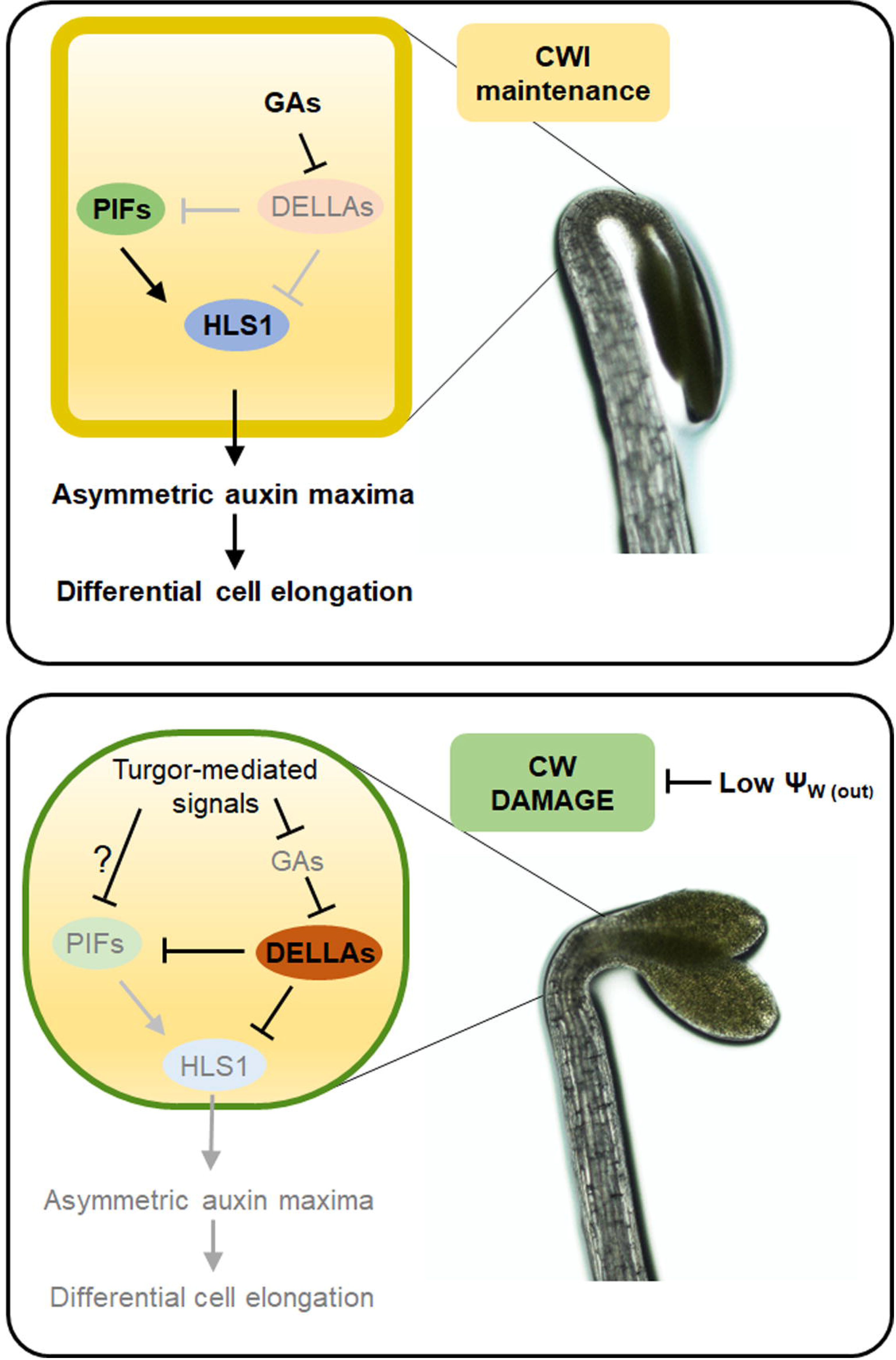
Proposed model of the effects of loss of cell wall integrity on apical hook formation. Perturbation of cell wall integrity (CWI), either caused by mutations in pectin composition or by isoxaben, activates turgor-dependent responses that repress accumulation of active gibberellins (GAs), leading to stabilization of DELLA proteins and reduction of PIF4/PIF5 (PIFs) protein levels. Increased DELLAs and reduced PIFs result in impaired *HLS1* expression, impairing proper formation of auxin response maxima and differential cell elongation, and ultimately inhibiting apical hook development.

## Discussion

### Cell wall alterations impair differential cell elongation during apical hook formation in a turgor-dependent manner

Differential cell elongation is widely used in plants to adapt growth and development to external and endogenous signals. This is exemplified by apical hook formation, which is largely dependent on the differential cell elongation on the opposite sides of the hypocotyl apex (Guzman and Ecker, 1990; Abbas *et al*., 2013). Cell elongation occurs through the controlled expansion of the cell wall, which results from the interplay between turgor pressure and cell wall elasticity and extensibility (Ray *et al*., 1972). It is therefore not surprising that cell wall composition has a major impact on hook formation, and that an extensive interplay occurs between cell walls and the hormonal networks controlling hook formation (Aryal *et al*., 2020; Jonsson *et al*., 2021). However, despite our considerable knowledge of the signalling pathways controlling hook development, little is known of how cell wall-derived signals interact with these pathways to modulate differential cell expansion and hook bending. Here we have shown that changes in CWI, either caused by mutations in genes affecting pectin composition or by interference with cellulose deposition triggered by isx, hinder hook formation in Arabidopsis seedlings in a turgor-dependent manner. Moreover, altered pectin integrity in *qua2* seedlings compromises, in a turgor-dependent manner, asymmetric auxin maxima formation and differential cell elongation. Additionally, we show that turgor-mediated responses triggered by altered CWI downregulate a hook-promoting signalling events that are positively regulated by GAs and include PIF4/5 accumulation and *HLS1* expression (Figure 8). These results suggest that turgor pressure links CWI to a GA-dependent signalling pathway to modulate hook formation and maintenance.

Cell wall assembly and remodelling must be finely controlled during growth processes to ensure proper cell expansion while maintaining mechanical integrity (Wolf *et al*., 2012). Moreover, alterations in CWI can occur in response to abiotic or biotic stress (Vaahtera *et al*., 2019; Lorrai *et al*., 2021); therefore, the structural and functional integrity of the wall must be constantly monitored and fine-tuned to allow normal growth and development under unstressed conditions while preventing mechanical failure under adverse conditions (Rui and Dinneny, 2020). Increasing evidence points to a role of turgor-mediated responses in triggering several effects of loss of CWI on plant growth and development (Verger *et al*., 2018; Engelsdorf *et al*., 2018). Indeed, plant cells must sustain huge turgor pressures, and their connection with each other, which is mediated by the cell wall, allows the propagation of signals generated by turgor pressure and by differential growth (Jonsson *et al*., 2022). Plants with altered CWI may fail to counterbalance turgor pressure, causing mechanical stress and triggering downstream compensatory responses. Indeed, supplementation with osmolytes, like sorbitol, or increasing medium agar concentrations has been previously exploited to decrease turgor pressure and restore growth in plants with perturbed cell walls (Verger *et al*., 2018; Engelsdorf *et al*., 2018; Bacete *et al*., 2022). We have shown here that both sorbitol and HA restore hook development in plants with altered pectin composition (Figure 2; Figure S2). Indeed, analysis of cell growth rate showed that the impaired hook formation phase observed in *qua2* is accompanied by a reduction of cell elongation rate in the outer cell layer (Figure 3a-b). Moreover, WT-like growth rate was restored when *qua2* seedlings were grown in HA condition (Figure 3a-b), further supporting the hypothesis that the defect in hook development observed in this mutant is largely mediated by turgor-dependent mechanisms.

It has been proposed that loss of cell adhesion in plants with altered HG is a consequence of excessive tension in the epidermis caused by mechanical stress (Verger *et al*., 2018). Moreover, tension-mediated signals triggered by altered pectin composition might induce compensatory mechanisms that restrict cell expansion and therefore relieve mechanical stress. We have previously observed that the reduced cell expansion observed in *qua2* seedlings is at least partly mediated by an increased expression of *AtPRX71*, encoding a ROS-generating apoplastic peroxidase (Raggi *et al*., 2015). Notably, *AtPRX71* expression is also induced by hypoosmolarity (Rouet *et al*., 2006), a condition leading to excessive turgor pressure. This suggests that turgor-dependent responses triggered by altered pectin composition might lead to compensatory mechanisms, possibly including peroxidase-mediated cell wall crosslinking, that ultimately restrict cell expansion. Such mechanisms might take place also during apical hook formation, causing the turgor-dependent defect in differential cell expansion observed in *qua2*.

The observation that isx also impairs hook formation under LA, but not HA conditions (Figure 6a) indicates that loss of CWI caused not only by pectin alterations, but also by defects in cellulose deposition trigger turgor-dependent signals that hinder differential cell expansion. However, the exact nature of the signals linking altered CWI to turgor-dependent suppression of cell expansion still needs to be clarified. It has been proposed that loss of CWI results in distortion or displacement of the plasma membrane relative to the cell wall, that can be detected by a dedicated CWI maintenance mechanism (Engelsdorf *et al*., 2018). Further investigation will provide insights in the role of specific components of the CWI maintenance system in modulating differential cell expansion during hook formation.

### Loss of CWI represses a signalling module that promotes apical hook development

Differential elongation during hook development requires the formation of an auxin gradient, reaching a maximum in the inner side of the hook where it reduces cell growth rate (Abbas *et al*., 2013). The cell wall is a key hub in this process, as a positive feedback loop mechanism couples cell wall stiffness, mediated by changes in the DM of HG with auxin redistribution (Jonsson *et al*., 2021). However, the mechanisms linking changes in cell wall properties and the signalling pathways that modulate differential cell expansion are poorly understood. Our results suggest that CWI represses a signalling module, comprising PIF4/5 and HLS1 (Figure 4 and 6), that positively regulates auxin biosynthesis and distribution and ultimately hook formation (Lehman *et al*., 1996; Franklin *et al*., 2011; Zhang *et al*., 2018). HLS1 suppresses accumulation of AUXIN RESPONSE FACTOR 2 (ARF2) (Li *et al*., 2004), which negatively regulates hook formation and transcriptional control of auxin transporters downstream of xyloglucan defects (Aryal *et al*., 2020). Our results indicate that mutants with altered pectin composition and seedlings treated with isx show a reduction of *HLS1* and *PIF4/5* transcript levels (Figure 4 and 6), and that isx significantly reduces PIF4/5 protein levels in WT plants (Figure 6). The downregulation of *HLS1* and *PIFs* might contribute to the disruption of asymmetric auxin maxima and of differential cell expansion observed in *qua2* and might also contribute to the hook defect caused by altered cellulose deposition, as HLS1 overexpression confers a partial resistance to the inhibitory effect of isx (Figure 6d), pointing to a common regulation of hook formation in response to changes in different cell wall components.

Mechanical stress arising from turgor pressure changes might activate JA-mediated stress responses in plants with altered CWI (Engelsdorf *et al*., 2018). Recently, it has been proposed that JA-Ile accumulation in the roots of the *kor1* mutant is prompted by turgor-driven mechanical compression at the level of the cortex (Mielke *et al*., 2021). We found that *qua2* seedlings accumulate high levels of jasmonates, which decrease when the mutant is grown in HA conditions (Figure 7a), confirming that cell wall stress-induced JA production is mediated by turgor pressure changes. However, JA signalling does not appear to be involved in the repression of hook development caused by loss of CWI neither in *qua2* nor in isx-treated seedlings (Figure 7b-d). On the other hand, our results suggest that hook defects in plants with an altered cell wall might be at least partially mediated by a reduction in GA levels, as 1) GA_4_ levels are reduced in *qua2* seedlings (Figure 5a); 2) both pectin-related mutants and isx-treated plants show altered expression of genes involved in the homeostasis of GAs (Figure 5b-c and Figure S3a-c); 3) exogenous GAs restore hook formation in pectin mutants and in isx-treated WT seedlings (Figure 5d and 6g); 4) isx stabilizes the DELLA protein RGA (Figure 6i-j), whose nuclear accumulation is negatively regulated by GAs (Silverstone *et al*., 2001), and 5) lack of all five *Arabidopsis* DELLA proteins reduces the impact of isx on hook formation (Figure 6h). Notably, growth of seedlings on HA, which restores normal hook formation in both *qua2* and isx-treated seedlings (Figure 2 and 6a), also restores GA_4_ accumulation in *qua2* seedling (Figure 5a), and increases *HLS1*, *PIF4/5* and GA biosynthetic gene expression in both *qua2* and isx-treated seedlings (Figure 4b-c, Figure 5b-c, Figure 6b-c, Figure S3a-c). Furthermore, HA conditions prevent the reduction of PIF4/5 protein levels and the increase of RGA levels in isx-treated WT seedlings (Figure 6e-f and i-j; Figure S4b-c). These results suggest a causal link between altered CWI, reduction of GA levels and suppression of GA-mediated signalling required for proper auxin signalling and differential cell expansion during hook formation and maintenance. Future work will provide further insights in the relative role of turgor-mediated responses to loss of CWI and of biochemical signals derived from specific cell wall structural components in modulating the signalling pathways that promote hook formation and other plant developmental processes that rely on differential cell elongation.

In conclusion, our results indicate that turgor-dependent responses link changes in CWI to the downregulation of a regulatory module, comprising GAs, PIFs and HLS1, that promotes asymmetric cell elongation and hypocotyl curvature during hook formation (Figure 8). However, it cannot be ruled out that additional mechanisms might contribute to compromise hook formation in plants with defective cell wall composition. Intriguingly, it was reported that short fragments of HG restore hook development in dark-grown mutants impaired in pectin composition (Sinclair *et al*., 2017), suggesting that, in WT plants, HG-derived fragments might act as signals that promote hook formation. Future research will help elucidate the mechanisms linking changes in the cell wall biochemical and physical properties occurring in response to internal and environmental cues to the signalling cascades that modulate differential cell growth during plant developmental programs.

## Experimental Procedures

### Plant lines

The *qua2-1* mutant was a gift of Gregory Mouille (INRA Centre de Versailles-Grignon), *coi1-1* and *jar1-1* mutant were a gift of Edward Farmer (Department of Plant Molecular Biology, University of Lausanne). The *mur1-1*, *mur4-1*, *mur7-1*, *prc1-1*, *kor1-1*, *gae1-1 gea6-1* double mutant, the pentuple *della* mutant (*gai-t6*, *rga-t2*, *rgl1-1*, *rgl2-1* and *rgl3-1*) and the transgenic line expressing pRGA:RGA-GFP were obtained by the Nottingham Arabidopsis Stock Centre. Transgenic lines *PIF4p:PIF4-HA pif4-301* and *PIF5p:PIF5-HA pif5-3* were a gift of Christian Fankhauser (University of Lausanne, Center for Integrative Genomics). The 35S::Myc-HLS1/*hls1-1* was a gift by Shangwei Zhong (Peking University).

The *qua2-1 coi1-1* and *qua2-1 jar1-1* double mutant lines were generated by crossing single mutants. Double homozygous lines were isolated based on the presence of cell adhesion defects in the hypocotyl and on primary root resistance to exogenous JA. *qua2-1 coi1-1* double homozygous mutants were crossed with a *qua2-1/qua2-1 coi1-1/COI1* sesquimutant, and experiments were performed on seedlings of the segregating progeny that were insensitive to JA in terms of root elongation.

The *qua2* DR5-VENUS line was generated by crossing a WT line expressing DR5-VENUS (*pDR5rev::3XVENUS-N7*) (Heisler *et al*., 2005) with *qua2-1*. The *qua2* myr-YFP line, expressing the myr-YFP plasma membrane marker line, was obtained by crossing a WT line carrying the pUBQ10::myr:YFP construct (Willis *et al*., 2016) with an homozygous *qua2-1* line. In both cases, double *qua2-1* homozygous lines were isolated based on the presence of cell adhesion defects in the hypocotyl, and homozygosity of the transgene was confirmed based on signal fluorescence in the F3 generation.

All lines used in this work were in the Col-0 background, except for *kor1-1*, in Wassilewskija (Ws) background, and *della*, in Landsberg *erecta* (L*er*) background.

### Plant growth conditions

Seeds were surface sterilised with absolute ethanol (v/v), air dried and sown on solid medium containing 2.2 gL^-1^ Murashige-Skoog (MS) salts (Duchefa), 1% (w/v) Suc, 0.8% or 2.5% (w/v) plant agar (Duchefa), pH 5.6. Plates were wrapped in aluminium foil and stratified at +4°C for 2-3 days. Isx (Merck) was dissolved in 0.01% dimethylsulphoxide (DMSO) and supplemented to growth medium at indicated concentrations. For etiolated growth, after stratification, germination was induced by exposure to white light for 4-6 hours, plates were wrapped in aluminium foils and placed in a growth chamber for indicated days. For hook angle analysis with sorbitol supplementation, seeds were sown on a sterilised nylon mesh placed on agar medium plates without sorbitol and placed in the dark as described above. After 24 h, the nylon mesh was transferred under green dim light to new plates containing sorbitol. All supplements were added in the indicated concentrations to autoclaved control media. For RNA and protein analysis, seedlings were harvested under dim green light and flash frozen in liquid nitrogen.

### Kinematic analysis of apical hook development and cell elongation measurement

Seedlings were grown vertically on solid medium plates in the dark at 21°C, illuminated with far infra-red light (850 nm). Seedlings were photographed every hour using a RASPBERRY PI camera. Apical hook angles were measured using Image J software (http://imagej.nih.gov/ij/).

For time-lapse imaging of cell expansion, WT myr-YFP and *qua2* myr-YFP seedlings were imaged using a Zeiss LSM800 confocal microscope equipped with 10x/0.45 Plan-apo dry objective. Z-stacks were acquired without averaging with a 0.5-micron cubic voxel size. Dark grown seedlings were placed on an agar gel block on a microscopy slide and imaged at three-hour intervals. Between acquisition of images, seedlings were placed vertically in a dark chamber to maintain skotomorphogenic conditions. Cell elongation was calculated using the software MorphographX (MGX). Using MGX, epidermal cell surface area from Z-stacks was extracted as described previously (Barbier de Reuille *et al*., 2015). Longitudinal expansion was calculated in MGX by overlaying Z-stacks with a fitted curved Bezier grid providing axial growth coordinates. For each condition and genotype, 15 cells from both the inner and the outer side of the hook were measured from each of 9 individual seedlings (135 cells). The data was statistically analysed by two-tailed Student’s t-test.

### Gene expression analysis

To analyse gene expression, uppermost part of seedling hypocotyls, including the apical hook, were isolated using a razor blade, frozen in liquid nitrogen and homogenised with an MM301 Ball Mill (Retsch) mixer ill for about 1 min at 25Hz. Total RNA was extracted with NucleoZOL reagent (Macherey-Nagel) according to the manufacturer’s instructions. 1μg of total RNA was retrotranscribed with Improm II Reverse Transcriptase (Promega). cDNA was mixed with iTaq Universal SYBR Green Supermix (Bio-Rad) and primer pairs specific for genes of interest, loaded onto 96-well plates and samples were amplified using a CFX96 Real-time System (Bio-Rad). Gene expression, relative to *UBIQUITIN5* (*UBQ5*), was calculated according to the ΔΔCT method.

### Protein extraction and immunoblot assays

Total proteins were extracted from etiolated seedlings (n=30) grounded in liquid nitrogen and resuspended in 120 µL of extraction buffer [125 mM Tris, pH 6.8, 4% (w/v) SDS, 20% (v/v) glycerol, 0.02% (w/v) bromophenol blue, 10% (v/v) β-mercaptoethanol]. Samples were heated for 5 min at 95°C and centrifuged for 1 min at 15,000g at room temperature. Proteins (20 µL of each sample) were separated by 8% acrylamide SDS-PAGE and transferred to a nitrocellulose membrane using the Trans-Blot Turbo transfer kit (Bio-Rad). 5% (w/v) milk dissolved in phosphate-buffered saline with 0.05% (v/v) Tween 20 was used for blocking for 1.5 h at room temperature and antibody dilutions. For the detection of HA, a 1:1000 dilution of the (F-7) sc-7392 antibody (Santacruz) was used. For the detection of GFP, a 1:1000 dilution of primary antibody (Chromotek) was used. As a secondary antibody, a 1:2000 dilution of a HRP-conjugated anti-mouse immunoglobulin (Cell Signaling) was used. Anti-actin polyclonal primary antibody (Agrisera) was used as loading control, with HRP-conjugated anti-rabbit immunoglobulin (1:2000; Cell Signaling) as a secondary antibody. The chemiluminescent signal of HRP conjugated to secondary antibodies was detected with ECL Western Blotting Substrate (Promega) using a ChemiDoc XRS+ system (Biorad).

### Confocal laser-scanning microscopy

For DR5::VENUS detection, three-day-old etiolated seedlings were placed between a microscopy slide and a cover slip. Images were made using a Zeiss LSM 880 laser scanning confocal microscope. Images were acquired using the Zen black software, with a 40X (C-Apochromat 40x/1.2 W Korr FCS M27) objective. Z-stacks were acquired without averaging with the image size 1024×1024 px and 0.345-micron pixel size and a Z-step size of 1μm. PI excitation was performed at 561 nm wavelength and the emission was collected in the range of 562-600 nm. VENUS excitation was performed at 514 nm wavelength and the emission was collected in the 518-560 nm range. The laser reflection was filtered by a beam splitter.

### Hormone quantification

Three-day-old dark-grown seedlings were homogenised with mortar and pestle in liquid nitrogen and reweighted into three replicates for gibberellin analysis and three replicates for jasmonates analysis (approximately 10 mg per sample).

Extraction and analysis of endogenous GA_4_ were performed according to the modified method described in Urbanová *et al*., (2013). Briefly, tissue samples of about 10 mg FW were ground to a fine consistency using 2.8-mm zirconium oxide beads (Retsch GmbH & Co. KG, Haan, Germany) and a MM 400 vibration mill at a frequency of 27 Hz for 3 min (Retsch GmbH & Co. KG, Haan, Germany) with 1 mL of ice-cold 80% acetonitrile containing 5% formic acid as extraction solution. The samples were then extracted overnight at 4°C using a benchtop laboratory rotator Stuart SB3 (Bibby Scientific Ltd., Staffordshire, UK) after adding internal standard ([^2^H_2_]GA_4_, OlChemIm, Czech Republic). The homogenates were centrifuged at 42030x*g* and 4°C for 10 min, and the corresponding supernatants were further purified using mixed-mode SPE cartridges (Waters, Milford, MA, USA). The samples were finally analysed with a UHPLC-MS/MS system consisting of the Acquity I-class UPLC^®^ chromatograph (Waters, Milford, MA, USA) coupled to the Xevo TQ-XS triple quadrupole mass spectrometer (Micromass, Manchester, UK). Masslynx 4.2 software (Waters, Milford, MA, USA) was used to analyse the acquired data.

Analysis of jasmonates was performed following a previously described protocol (Floková *et al*., 2014). Briefly, the samples were extracted in 1 mL of ice-cold 10% aqueous methanol with the addition of isotopically labelled internal standards ([^2^H_6_]-JA and [^2^H_2_]–(–)– JA-Ile, purchased from OlChemIm, Czech Republic) and the resulting extracts were purified on Oasis® HLB SPE columns (1 cc/30 mg, Waters). The analyses were carried out using a 1290 Infinity liquid chromatography system coupled to an Agilent 6490 Triple Quadrupole mass spectrometer (Agilent Technologies, Santa Clara, CA, USA). The data were processed in MassHunter Quantitative B.09.00 software (Agilent Technologies, Santa Clara, CA, USA) (Agilent Technologies, Santa Clara, CA, USA) (Široká *et al*., 2022).

### Accession numbers

AT1G78240 (*QUA2*); AT4G37580 (*HLS1*); AT2G43010 (*PIF4*); AT3G59060 (*PIF5*); AT1G15550 (*GA3ox1*); AT4G25420 (*GA20ox1*); AT1G30040 (*GA2ox2*); AT3G51160 (*MUR1*); AT1G30620 (*MUR4*); AT4G30440 (GAE1); AT3G23820 (*GAE6*); AT5G64740 (PRC1); AT5G49720 (*KOR1*); AT2G46370 (*JAR1*); AT2G39940 (*COI1*); AT2G01570 (*RGA*); AT1G14920 (*GAI*); AT1G66350 (*RGL1*); AT3G03450 (*RGL2*); AT5G17490 (*RGL3*).

## Acknowledgements

We are grateful to Christian Fankhauser (University of Lausanne) for providing *PIF4p:PIF4-HApif4-301* and *PIF5p:PIF5-HApif5-3* seeds, to Edward Farmer (University of Lausanne) for providing *coi1-1* and *jar1-1* seeds, to Shangwei Zhong (Peking University) for providing 35S:Myc-HLS1/*hls1-1* seeds. The authors acknowledge the facilities and technical assistance of the Umeå Plant Science Centre (UPSC) Microscopy facility.

This work was supported by Sapienza University of Rome (“Progetti di Avvio alla Ricerca 2021 – Tipo 2” grant n. AR22117A5E76C7EE, awarded to R.L.; “Progetti di Ricerca 2021 – Progetti Medi” grant n. RM12218161B8A750 and Progetti di Ricerca 2019 “Progetti Medi” grant n. RM11916B6F156C03, awarded to S.F.) and by Regione Lazio (grant n. A0375-2020-36720 “Alternative use of agri-food waste in a circular economy context”, call Lazioinnova for Research Group Projects 2020, awarded to S.F.). This work was also supported by grants from the Knut and Alice Wallenberg Foundation (KAW 2016.0341, KAW 2016.0352 and KAW 2022.0029 (SaR)), the Swedish Governmental Agency for Innovation Systems (VINNOVA 2016-00504) and Vetenskapsrådet VR-2020-03420 (SaR, S.R.). We also thank Bio4Energy, a Strategic Research Environment supported through the Swedish Government’s Strategic Research Area initiative, for supporting this work. R.L. is supported by the Italian Ministry of University and Research (MUR) (project “Development of bio-based solutions for the valorisation of waste agri-food biomass”, D.M. n. 1062-10.08.2021 PON “Ricerca e Innovazione” 2014-2020, Asse IV “Istruzione e ricerca per il recupero” – Azione IV.4 – “Dottorati e contratti di ricerca su tematiche dell’innovazione” e Azione IV.6 – “Contratti di ricerca su tematiche Green”). J.S. was financially supported by Czech Science Foundation project No. 19-10464Y. J.S. and O.N. thank Miroslava Špičáková for her technical support. D.T. is grateful for technical assistance of Renata Plotzova and for financial support from the Ministry of Education, Youth and Sports of the Czech Republic (European Regional Development Fund-Project ‘Centre for Experimental Plant Biology’ no. CZ.02.1.01/0.0/0.0/16_019/0000738).

This study was carried out within the Agritech National Research Center and received funding from the European Union Next-GenerationEU (PIANO NAZIONALE DI RIPRESA E RESILIENZA (PNRR) – MISSIONE 4 COMPONENTE 2, INVESTIMENTO 1.4 – D.D. 1032 17/06/2022, CN00000022). This manuscript reflects only the authors’ views and opinions, neither the European Union nor the European Commission can be considered responsible for them.

## Supplementary materials

**Figure S1. Apical hook angle in pectin mutants grown on low and high agar.** Wild-type (WT), *qua2*, *mur1* and *gae1gae6* seedlings were grown for three days in the dark on medium containing either 0.8% (w/v) (LA, white boxes) or 2.5% (w/v) agar (HA, orange boxes). Box plots indicate the 1^st^ and 3^rd^ quartiles split by median; whiskers show range. Letters indicate statistically significant differences (P < 0.05) according to two-way ANOVA followed by post-hoc Tukey’s HSD.

**Figure S2. Osmotic support suppresses apical hook defects in pectin mutants.** Wild-type (WT), *qua2*, *mur1* and *gae1 gae6* seeds were germinated in the dark on solid medium without sorbitol and, 24 h after germination, were transferred to solid medium containing increasing sorbitol concentrations (white boxes, 0 mM; yellow boxes, 125 mM; red boxes, 250 mM). After three days from light exposure, the apical hook angle was measured. Box plots indicate the 1^st^ and 3^rd^ quartiles split by median; whiskers show range (n>20). Letters indicate statistically significant differences (P < 0.05) according to two-way ANOVA followed by post-hoc Tukey’s HSD.

**Figure S3. Effects of isoxaben on the expression of genes involved in GA metabolism.** Expression levels of *GA3ox1* (**a**), *GA20ox1* (**b**) and *GA2ox2* (**c**) were determined in WT seedlings grown in the dark for three dayes on medium containing 0.8% (LA) or 2.5% (HA) agar (w/v) in the presence of DMSO (mock) or 2.5 nM isoxaben (isx). Transcript levels were determined by qPCR using *UBQ5* as reference gene. Bars indicate mean relative expression levels ± SD (n≥3), compared to WT seedlings grown in LA in the absence of isx. Asterisks indicate statistically significant differences with WT in LA conditions, according to Student’s t-test (*, p≤0.05; **, p≤0.01).

**Figure S4. Effects of isoxaben on PIF5 expression.** (**a**), WT seedlings were grown in the dark in the presence of DMSO (mock) or 2.5 nM isoxaben (isx) on medium containing 0.8% (LA) or 2.5% (HA) agar (w/v). *PIF5* transcript levels were determined by qPCR using *UBQ5* as reference. Bars indicate mean relative expression, compared to WT seedlings grown in LA in the absence of isx, ± SD of at least three independent biological replicates. Asterisks indicate statistically significant differences with WT in LA conditions, according to Student’s t-test (*, p≤0.05). **(b**), Transgenic lines expressing PIF5-HA under the control of its native promoter (ProPIF5:PIF5-3×HA) were germinated and grown in the dark for three days in LA or HA medium supplemented with DMSO (mock), 2.5 nM isx or 5 nM isx. PIF5-HA levels were detected by immunoblot analysis using an antibody against HA. An antibody against Actin (ACT) was used as loading control. (**c**), Intensity of the PIF5-HA signal in (**b**), normalized for the ACT signal, was quantified and expressed as relative levels compared to mock-treated seedlings under LA condition. (c). Bars indicate means of n≥3 independent biological replicates ± SD. Asterisks indicate statistically significant differences between mock-treated and isx-treated seedlings in the same agar concentrations, according to Student’s t test (*P < 0.05).

**Table S1**. Primers used for qRT-PCR analysis and genotyping.

## References

1. Abbas, M., Alabadí, D. and Blázquez, M. (2013) Differential growth at the apical hook: all roads lead to auxin. Frontiers in Plant Science, 4. Available at: https://www.frontiersin.org/articles/10.3389/fpls.2013.00441 [Accessed November 9, 2022].

2. An, F., Zhang, X., Zhu, Z., Ji, Y., He, W., Jiang, Z., Li, M. and Guo, H. (2012) Coordinated regulation of apical hook development by gibberellins and ethylene in etiolated Arabidopsis seedlings. Cell research, 22, 915–927.

3. Aryal, B., Jonsson, K., Baral, A., Sancho-Andres, G., Routier-Kierzkowska, A.-L., Kierzkowski, D. and Bhalerao, R.P. (2020) Interplay between Cell Wall and Auxin Mediates the Control of Differential Cell Elongation during Apical Hook Development. Current Biology, 30, 1733–1739.e3.

4. Bacete, L., Schulz, J., Engelsdorf, T., et al. (2022) THESEUS1 modulates cell wall stiffness and abscisic acid production in Arabidopsis thaliana. Proc Natl Acad Sci U S A, 119, e2119258119.

5. Baral, A., Aryal, B., Jonsson, K., et al. (2021) External Mechanical Cues Reveal a Katanin-Independent Mechanism behind Auxin-Mediated Tissue Bending in Plants. Developmental Cell, 56, 67–80.e3.

6. Barbier de Reuille, P., Routier-Kierzkowska, A.-L., Kierzkowski, D., et al. (2015) MorphoGraphX: A platform for quantifying morphogenesis in 4D D. C. Bergmann, ed. eLife, 4, e05864.

7. Bethke, G., Thao, A., Xiong, G., et al. (2016) Pectin Biosynthesis Is Critical for Cell Wall Integrity and Immunity in Arabidopsis thaliana. The Plant Cell, 28, 537–556.

8. Bonin, C.P., Potter, I., Vanzin, G.F. and Reiter, W.-D. (1997) The MUR1 gene of Arabidopsis thaliana encodes an isoform of GDP-d-mannose-4,6-dehydratase, catalyzing the first step in the de novo synthesis□of□GDP-l-fucose. Proceedings of the National Academy of Sciences, 94, 2085–2090.

9. Bouton, S., Leboeuf, E., Mouille, G., Leydecker, M.-T., Talbotec, J., Granier, F., Lahaye, M., Höfte, H. and Truong, H.-N. (2002) QUASIMODO1 Encodes a Putative Membrane-Bound Glycosyltransferase Required for Normal Pectin Synthesis and Cell Adhesion in Arabidopsis. The Plant Cell, 14, 2577–2590.

10. Burget, E.G., Verma, R., Mølhøj, M. and Reiter, W.-D. (2003) The Biosynthesis of l-Arabinose in Plants: Molecular Cloning and Characterization of a Golgi-Localized UDP-d-Xylose 4-Epimerase Encoded by the MUR4 Gene of Arabidopsis. The Plant Cell, 15, 523–531.

11. Cosgrove, D.J. (2005) Growth of the plant cell wall. Nature reviews molecular cell biology, 6, 850–861.

12. De Lucas, M., Daviere, J.-M., Rodríguez-Falcón, M., et al. (2008) A molecular framework for light and gibberellin control of cell elongation. Nature, 451, 480–484.

13. Desnos, T., Orbovic, V., Bellini, C., Kronenberger, J., Caboche, M., Traas, J. and Hofte, H. (1996) Procuste1 mutants identify two distinct genetic pathways controlling hypocotyl cell elongation, respectively in dark– and light-grown Arabidopsis seedlings. Development, 122, 683–693.

14. Du, J., Kirui, A., Huang, S., et al. (2020) Mutations in the Pectin Methyltransferase QUASIMODO2 Influence Cellulose Biosynthesis and Wall Integrity in Arabidopsis. The Plant Cell, 32, 3576–3597.

15. Ellis, C., Karafyllidis, I., Wasternack, C. and Turner, J.G. (2002) The Arabidopsis Mutant cev1 Links Cell Wall Signaling to Jasmonate and Ethylene Responses. The Plant Cell, 14, 1557–1566.

16. Engelsdorf, T., Gigli-Bisceglia, N., Veerabagu, M., McKenna, J.F., Vaahtera, L., Augstein, F., Van der Does, D., Zipfel, C. and Hamann, T. (2018) The plant cell wall integrity maintenance and immune signaling systems cooperate to control stress responses in Arabidopsis thaliana. Science Signaling, 11, eaao3070.

17. Fagard, M., Desnos, T., Desprez, T., et al. (2000) PROCUSTE1 Encodes a Cellulose Synthase Required for Normal Cell Elongation Specifically in Roots and Dark-Grown Hypocotyls of Arabidopsis. The Plant Cell, 12, 2409–2423.

18. Feng, S., Martinez, C., Gusmaroli, G., et al. (2008) Coordinated regulation of Arabidopsis thaliana development by light and gibberellins. Nature, 451, 475–479.

19. Floková, K., Tarkowská, D., Miersch, O., Strnad, M., Wasternack, C. and Novák, O. (2014) UHPLC–MS/MS based target profiling of stress-induced phytohormones. Phytochemistry, 105, 147–157.

20. Franklin, K.A., Lee, S.H., Patel, D., et al. (2011) PHYTOCHROME-INTERACTING FACTOR 4 (PIF4) regulates auxin biosynthesis at high temperature. Proceedings of the National Academy of Sciences, 108, 20231–20235.

21. Freshour, G., Bonin, C.P., Reiter, W.-D., Albersheim, P., Darvill, A.G. and Hahn, M.G. (2003) Distribution of Fucose-Containing Xyloglucans in Cell Walls of the mur1 Mutant of Arabidopsis. Plant Physiology, 131, 1602.

22. Guzman, P. and Ecker, J.R. (1990) Exploiting the triple response of Arabidopsis to identify ethylene-related mutants. The Plant Cell, 2, 513–523.

23. Hamann, T., Bennett, M., Mansfield, J. and Somerville, C. (2009) Identification of cell-wall stress as a hexose-dependent and osmosensitive regulator of plant responses. The Plant Journal, 57, 1015–1026.

24. Hedden, P. and Phillips, A.L. (2000) Gibberellin metabolism: new insights revealed by the genes. Trends in Plant Science, 5, 523–530.

25. Heisler, M.G., Ohno, C., Das, P., Sieber, P., Reddy, G.V., Long, J.A. and Meyerowitz, E.M. (2005) Patterns of Auxin Transport and Gene Expression during Primordium Development Revealed by Live Imaging of the Arabidopsis Inflorescence Meristem. Current Biology, 15, 1899–1911.

26. Jonsson, K., Hamant, O. and Bhalerao, R.P. (2022) Plant cell walls as mechanical signaling hubs for morphogenesis. Current Biology, 32, R334–R340.

27. Jonsson, K., Lathe, R.S., Kierzkowski, D., Routier-Kierzkowska, A.-L., Hamant, O. and Bhalerao, R.P. (2021) Mechanochemical feedback mediates tissue bending required for seedling emergence. Current Biology, 31, 1154–1164.e3.

28. Kaczmarska, A., Pieczywek, P.M., Cybulska, J. and Zdunek, A. (2022) Structure and functionality of Rhamnogalacturonan I in the cell wall and in solution: A review. Carbohydrate Polymers, 278, 118909.

29. Khanna, R., Shen, Y., Marion, C.M., Tsuchisaka, A., Theologis, A., Schäfer, E. and Quail, P.H. (2007) The Basic Helix-Loop-Helix Transcription Factor PIF5 Acts on Ethylene Biosynthesis and Phytochrome Signaling by Distinct Mechanisms. Plant Cell, 19, 3915–3929.

30. Krupková, E., Immerzeel, P., Pauly, M. and Schmülling, T. (2007) The TUMOROUS SHOOT DEVELOPMENT2 gene of Arabidopsis encoding a putative methyltransferase is required for cell adhesion and co-ordinated plant development. The Plant Journal, 50, 735–750.

31. Lehman, A., Black, R. and Ecker, J.R. (1996) HOOKLESS1, an ethylene response gene, is required for differential cell elongation in the Arabidopsis hypocotyl. Cell, 85, 183– 194.

32. Li, H., Johnson, P., Stepanova, A., Alonso, J.M. and Ecker, J.R. (2004) Convergence of signaling pathways in the control of differential cell growth in Arabidopsis. Developmental cell, 7, 193–204.

33. Li, K., Yu, R., Fan, L.-M., Wei, N., Chen, H. and Deng, X.W. (2016) DELLA-mediated PIF degradation contributes to coordination of light and gibberellin signalling in Arabidopsis. Nature communications, 7, 1–11.

34. Lorrai, R., Francocci, F., Gully, K., Martens, H.J., De Lorenzo, G., Nawrath, C. and Ferrari, S. (2021) Impaired Cuticle Functionality and Robust Resistance to Botrytis cinerea in Arabidopsis thaliana Plants With Altered Homogalacturonan Integrity Are Dependent on the Class III Peroxidase AtPRX71. Front Plant Sci, 12, 696955.

35. Mielke, S., Zimmer, M., Meena, M.K., Dreos, R., Stellmach, H., Hause, B., Voiniciuc, C. and Gasperini, D. (2021) Jasmonate biosynthesis arising from altered cell walls is prompted by turgor-driven mechanical compression. Science Advances, 7, eabf0356.

36. Mølhøj, M., Verma, R. and Reiter, W.-D. (2004) The Biosynthesis of d-Galacturonate in Plants. Functional Cloning and Characterization of a Membrane-Anchored UDP-d-Glucuronate 4-Epimerase from Arabidopsis. Plant Physiology, 135, 1221–1230.

37. Mouille, G., Ralet, M.-C., Cavelier, C., et al. (2007) Homogalacturonan synthesis in Arabidopsis thaliana requires a Golgi-localized protein with a putative methyltransferase domain. The Plant Journal, 50, 605–614.

38. Nicol, F., His, I., Jauneau, A., Vernhettes, S., Canut, H. and Höfte, H. (1998) A plasma membrane-bound putative endo-1,4-β-D-glucanase is required for normal wall assembly and cell elongation in Arabidopsis. The EMBO Journal, 17, 5563–5576.

39. Raggi, S., Ferrarini, A., Delledonne, M., Dunand, C., Ranocha, P., De Lorenzo, G., Cervone, F. and Ferrari, S. (2015) The Arabidopsis Class III Peroxidase AtPRX71 Negatively Regulates Growth under Physiological Conditions and in Response to Cell Wall Damage. Plant Physiology, 169, 2513–2525.

40. Ray, P.M., Green, P.B. and Cleland, R. (1972) Role of turgor in plant cell growth. Nature, 239, 163–164.

41. Rayon, C., Cabanes-Macheteau, M., Loutelier-Bourhis, C., Salliot-Maire, I., Lemoine, J., Reiter, W.-D., Lerouge, P. and Faye, L. (1999) Characterization of N-Glycans from Arabidopsis. Application to a Fucose-Deficient Mutant1. Plant Physiology, 119, 725–734.

42. Reiter, W.-D., Chapple, C. and Somerville, C.R. (1997) Mutants of Arabidopsis thaliana with altered cell wall polysaccharide composition. The Plant Journal, 12, 335–345.

43. Reiter, W.-D., Chapple, C.C.S. and Somerville, C.R. (1993) Altered Growth and Cell Walls in a Fucose-Deficient Mutant of Arabidopsis. Science, 261, 1032–1035.

44. Rouet, M.-A., Mathieu, Y., Barbier-Brygoo, H. and Laurière, C. (2006) Characterization of active oxygen-producing proteins in response to hypo-osmolarity in tobacco and Arabidopsis cell suspensions: identification of a cell wall peroxidase. Journal of Experimental Botany, 57, 1323–1332.

45. Rui, Y. and Dinneny, J.R. (2020) A wall with integrity: surveillance and maintenance of the plant cell wall under stress. New Phytologist, 225, 1428–1439.

46. Shen, X., Li, Y., Pan, Y. and Zhong, S. (2016) Activation of HLS1 by Mechanical Stress via Ethylene-Stabilized EIN3 Is Crucial for Seedling Soil Emergence. Frontiers in Plant Science, 7. Available at: https://www.frontiersin.org/articles/10.3389/fpls.2016.01571 [Accessed July 12, 2023].

47. Silverstone, A.L., Jung, H.-S., Dill, A., Kawaide, H., Kamiya, Y. and Sun, T. (2001) Repressing a Repressor: Gibberellin-Induced Rapid Reduction of the RGA Protein in Arabidopsis. The Plant Cell, 13, 1555–1566.

48. Sinclair, S.A., Larue, C., Bonk, L., et al. (2017) Etiolated Seedling Development Requires Repression of Photomorphogenesis by a Small Cell-Wall-Derived Dark Signal. Current Biology, 27, 3403–3418.e7.

49. Široká, J., Brunoni, F., Pěnčík, A., Mik, V., Žukauskaitė, A., Strnad, M., Novák, O. and Floková, K. (2022) High-throughput interspecies profiling of acidic plant hormones using miniaturised sample processing. Plant Methods, 18, 122.

50. Song, S., Huang, H., Gao, H., et al. (2014) Interaction between MYC2 and ETHYLENE INSENSITIVE3 modulates antagonism between jasmonate and ethylene signaling in Arabidopsis. The Plant Cell, 26, 263–279.

51. Sun, T. (2008) Gibberellin metabolism, perception and signaling pathways in Arabidopsis. The Arabidopsis Book/American Society of Plant Biologists, 6.

52. Urbanová, T., Tarkowská, D., Novák, O., Hedden, P. and Strnad, M. (2013) Analysis of gibberellins as free acids by ultra performance liquid chromatography-tandem mass spectrometry. Talanta, 112, 85–94.

53. Vaahtera, L., Schulz, J. and Hamann, T. (2019) Cell wall integrity maintenance during plant development and interaction with the environment. Nature plants, 5, 924–932.

54. Verger, S., Long, Y., Boudaoud, A. and Hamant, O. (2018) A tension-adhesion feedback loop in plant epidermis C. S. Hardtke and D. C. Bergmann, eds. eLife, 7, e34460.

55. Wasternack, C. and Hause, B. (2013) Jasmonates: biosynthesis, perception, signal transduction and action in plant stress response, growth and development. An update to the 2007 review in Annals of Botany. Annals of Botany, 111, 1021–1058.

56. Willis, L., Refahi, Y., Wightman, R., Landrein, B., Teles, J., Huang, K.C., Meyerowitz, E.M. and Jönsson, H. (2016) Cell size and growth regulation in the Arabidopsis thaliana apical stem cell niche. Proceedings of the National Academy of Sciences, 113, E8238–E8246.

57. Wit, M. de, Keuskamp, D.H., Bongers, F.J., Hornitschek, P., Gommers, C.M.M., Reinen, E., Martínez-Cerón, C., Fankhauser, C. and Pierik, R. (2016) Integration of Phytochrome and Cryptochrome Signals Determines Plant Growth during Competition for Light. Current Biology, 26, 3320–3326.

58. Wolf, S., Hématy, K. and Höfte, H. (2012) Growth control and cell wall signaling in plants. Annual review of plant biology, 63, 381–407.

59. Yamaguchi, S. (2006) Gibberellin Biosynthesis in Arabidopsis. Phytochem Rev, 5, 39–47.

60. Zhang, B., Holmlund, M., Lorrain, S., Norberg, M., Bakó, L., Fankhauser, C. and Nilsson, O. (2017) BLADE-ON-PETIOLE proteins act in an E3 ubiquitin ligase complex to regulate PHYTOCHROME INTERACTING FACTOR 4 abundance H. Yu, ed. eLife, 6, e26759.

61. Zhang, X., Ji, Y., Xue, C., et al. (2018) Integrated regulation of apical hook development by transcriptional coupling of EIN3/EIL1 and PIFs in Arabidopsis. The Plant Cell, 30, 1971–1988.

62. Zhang, X., Zhu, Z., An, F., Hao, D., Li, P., Song, J., Yi, C. and Guo, H. (2014) Jasmonate-activated MYC2 represses ETHYLENE INSENSITIVE3 activity to antagonize ethylene-promoted apical hook formation in Arabidopsis. The Plant Cell, 26, 1105–1117.

