## Supplementary figures and images for "Cell wall integrity modulates a PHYTOCHROME-INTERACTING FACTOR (PIF) – HOOKLESS1 (HLS1) signalling module controlling apical hook formation in Arabidopsis"

### Figure S1

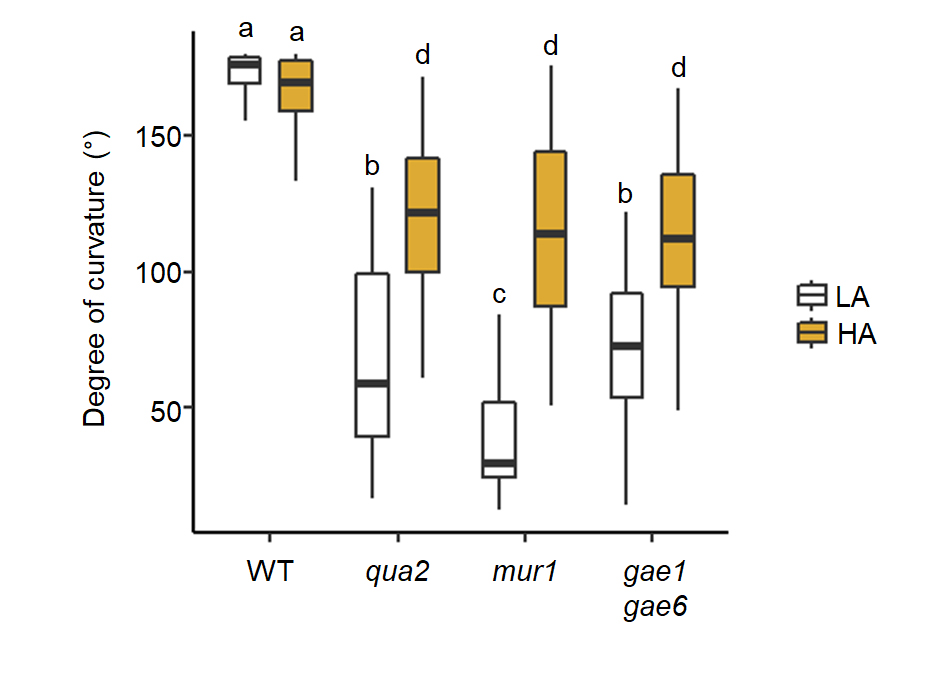

### Figure S2

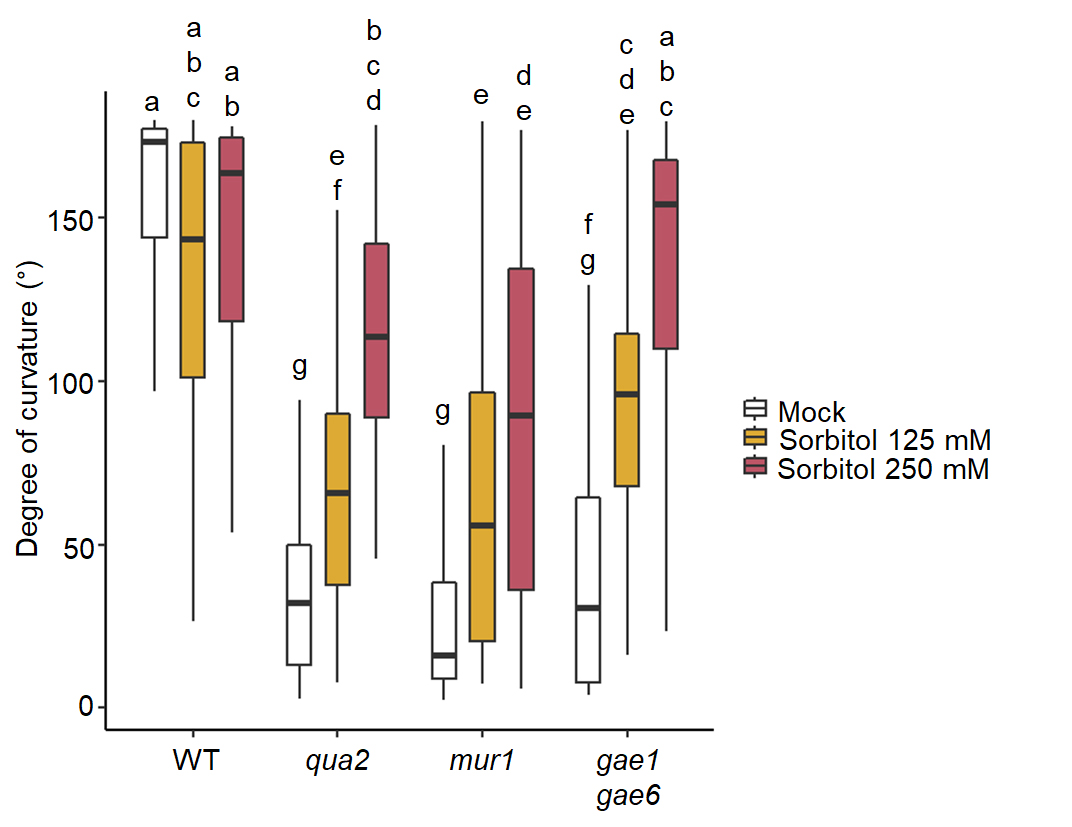

### Figure S3

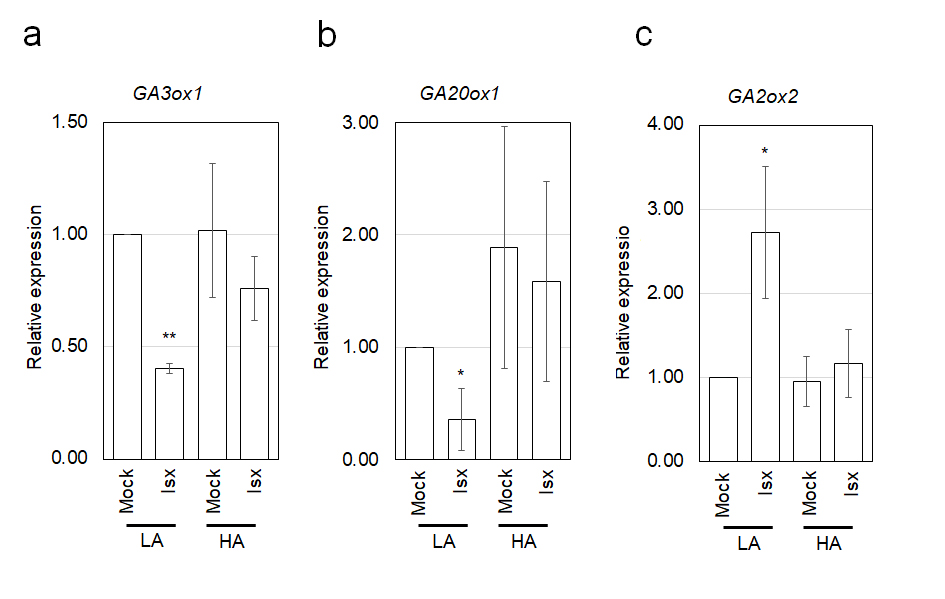

### Figure S4

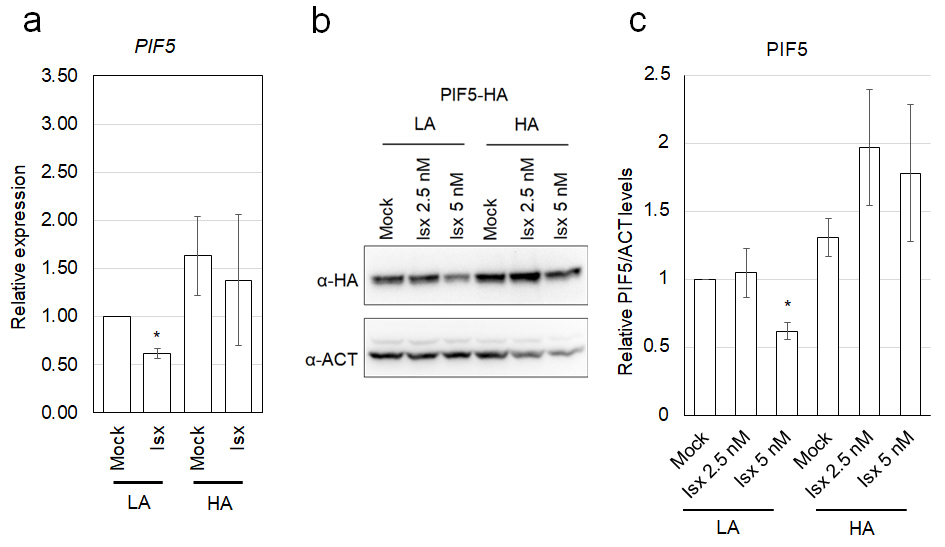
