## Supplementary material for "Cell wall integrity modulates a PHYTOCHROME-INTERACTING FACTOR (PIF) – HOOKLESS1 (HLS1) signalling module controlling apical hook formation in Arabidopsis": Table S1

| Gene | Sequence |
| --- | --- |
| *UBQ5* | GGAAGAAGAAGACTTACACC |
|  | AGTCCACACTTACCACAGTA |
| *PIF4* | TACCTCGATTTCCGGTTATGGATC |
|  | GTTGTTGACTTTGCTGTCCCGC |
| *PIF5* | GAGCAGCTCGCTAGGTACATG |
|  | GTTGTTGTTGCACGGTCTG |
| *GA3ox1* | GCTTAAGTCTGCTCGGTCGG |
|  | AGTGCGATACGAGCGACG |
| *GA20ox1* | AGCGAGAGGAAATCACTTGC |
|  | CGGCCCGGTTTTTAAGAGAC |
| *GA2ox2* | TCCGACCCGAACTCATGACT |
|  | CGGCCCGGTTTTTAAGAGAC |
| *COI1* | TGAAAGCATAGGCACATATCTGAA |
|  | ATTCACCTACGTAACCCAGCAGGAT |
| *HLS1* | GAATCCGACATTCACCTTCC |
|  | CATCCTCTAATCATGCCCACT |

**Supplemental Table S1.** The primer sequences used in this study.
